# Protein-lipid interactions drive presynaptic assembly upstream of cell adhesion molecules

**DOI:** 10.1101/2023.11.17.567618

**Authors:** Elisa B. Frankel, Araven Tiroumalechetty, Parise S. Henry, Zhaoqian Su, Yinghao Wu, Peri T. Kurshan

## Abstract

Textbook models of synaptogenesis position cell adhesion molecules such as neurexin as initiators of synapse assembly. Here we discover a mechanism for presynaptic assembly that occurs prior to neurexin recruitment, while supporting a role for neurexin in synapse maintenance. We find that the cytosolic active zone scaffold SYD-1 interacts with membrane phospholipids to promote active zone protein clustering at the plasma membrane, and subsequently recruits neurexin to stabilize those clusters. Employing molecular dynamics simulations to model intrinsic interactions between SYD-1 and lipid bilayers followed by *in vivo* tests of these predictions, we find that PIP_2_-interacting residues in SYD-1’s C2 and PDZ domains are redundantly necessary for proper active zone assembly. Finally, we propose that the uncharacterized yet evolutionarily conserved short γ isoform of neurexin represents a minimal neurexin sequence that can stabilize previously assembled presynaptic clusters, potentially a core function of this critical protein.

## INTRODUCTION

The orchestration of synapse assembly is fundamental to nervous system function, but many details of this process remain poorly understood. The canonical model of synapse formation posits that synaptic cell adhesion molecules (CAMs) recognize and bind specific trans-synaptic partners, an interaction which then initiates the recruitment of additional pre- and postsynaptic components^1–3^. Neurexins are a well-studied family of presynaptic CAMs that have been classified as central players in the initiation of synapse assembly based on their behaviors in artificial synapse formation assays^4–8^. However, the initiation of synapse assembly by neurexins or other CAMs has not been substantiated *in vivo,* leading to renewed questions about both the function of neurexin and, more generally, the mechanism of synapse assembly^4,5^.

In mammals, neurexins are encoded by three genes and expressed as α/long, β/medium, and γ/short isoforms, which differ in the length and complexity of their extracellular domains. The extracellular interactions of α- and β-neurexins have been studied extensively and likely underlie the specification of functionally diverse synaptic connections in mammals through extensive alternative splicing combined with interactions with numerous postsynaptic ligands^6–10^. Conditional knockouts of all α- and β-neurexins in mice led to defects in Ca^2+^ influx and synaptic strength across neuron types rather than widespread synapse loss^4^, suggesting that neurexins may serve primarily to specify synaptic functional attributes rather than acting as general initiators of synapse formation^5,11^. However, little is known about the function of the recently discovered γ/short-neurexin isoform^12–14^, which was not deleted in the conditional knockouts but lacks the structured extracellular domains that would conceivably mediate synapse-initiating trans-synaptic interactions.

In the simpler genetic paradigm of the invertebrate *C. elegans,* a single neurexin gene, *nrx-1*, encodes both α/long and γ/short neurexin isoforms^15,16^. In a strain containing presynaptic markers expressed in a single motor neuron, DA9, the complete deletion of *nrx-1* resulted in synapse loss that was limited to a region of the neuron receiving synapse elimination signals, suggesting that the function of NRX-1 is to stabilize and maintain synapses in the face of synapse-elimination cues, rather than to initiate their assembly^16^. Additionally, deletion of the α/long isoform of neurexin that preserved the γ/short isoform did not result in any loss or disruption of DA9 presynapses, indicating that the γ/short isoform can stabilize synapses even in the absence of canonical trans-synaptic interactions^16^. The persistence of the γ/short isoform of neurexin over hundreds of millions of years of divergent evolution suggests that it plays a critical function, but what that function is, whether and how it localizes to presynapses, and how widely γ/short neurexin is expressed in the nervous system are all unknown. Moreover, how synaptic components are initially assembled, if not through a neurexin-based trans-synaptic interaction mechanism, remains an important open question.

In both vertebrates and invertebrates, the cytosolic active zone scaffold proteins SYD-1 and SYD-2/Liprin-α have been identified as central regulators of presynaptic assembly^17–27^. In *C. elegans* and *D. melanogaster,* SYD-1 and SYD-2/Liprin-α arrive very early to nascent presynapses and function to recruit other active zone proteins^17–23,28^. *D. melanogaster* neurexin has been shown to interact with dSyd-1 through a PDZ-based interaction^29^, but the directionality of the relationship between these proteins, the mechanisms and interactions that drive their synaptic localization and clustering, and their joint contributions to synapse assembly and maintenance all have yet to be established^34^.

Here, we demonstrate that the initial clustering of endogenous presynaptic proteins *in vivo*, including SYD-1 and SYD-2, precedes the arrival of neurexin and instead depends on interactions with membrane phospholipids. We define an intracellular pathway directed by SYD-1 and mediated by interactions with plasma membrane PI(4,5)P_2_ that subsequently recruits and clusters neurexin through a PDZ-based interaction. Using molecular dynamics simulations followed by *in vivo* validation, we uncover the domains of SYD-1 that enable it to interact with PIP_2_ at the plasma membrane and accumulate at nascent presynaptic specializations before neurexin recruitment. Finally, we demonstrate that neurexin’s role in synapse stabilization is independent of extracellular interactions, suggesting a role for the evolutionarily conserved γ/short isoform.

Collectively, our studies establish the heretofore unappreciated central role of intracellular scaffold proteins in initiating presynaptic assembly through interactions with phospholipids in the cell membrane, independently of CAM-mediated trans-synaptic activation.

## RESULTS

### Neurexin stabilizes presynapses in the absence of its extracellular domain

*C. elegans* express two major neurexin isoform classes, long (α) and short (γ), which are driven by distinct promoters (Fig. 1A). The canonical transsynaptic binding domains of the α/long isoforms are absent in the γ/short isoforms of neurexin (Fig. 1B)^21^. To understand the roles of NRX-1 isoforms in synapse development *in vivo*, we assayed synapse formation in a single neuron. DA9 is a cholinergic motor neuron in the *C. elegans* tail, which lays out a set of *en passant* presynaptic specializations on the dorsal side of the animal at stereotypical locations along its axon (Fig. 1C). We previously showed that *nrx-1(-)* null mutants are characterized by loss of presynaptic active zones (labeled with active zone marker Clarinet(CLA-1)::GFP, the homolog of vertebrate Piccolo^30^), primarily in the most posterior region of the DA9 synaptic domain (Fig. 1D and ^16^, see Supp. Fig. 1A for examples of CLA-1::GFP puncta detection). Interestingly, we found that this spatially restricted phenotype occurs because the loss of *nrx-1* sensitizes presynaptic specializations of DA9 to elimination cues that are secreted from the posterior side. Thus, removal of the elimination signal, or the DA9-specific expression of either α/long or γ/short isoforms of NRX-1 can rescue this phenotype^16^ (Fig. 1D,E). This demonstrated that NRX-1(γ) isoforms are sufficient to stabilize active zones and prevent their elimination.

**Figure 1.**
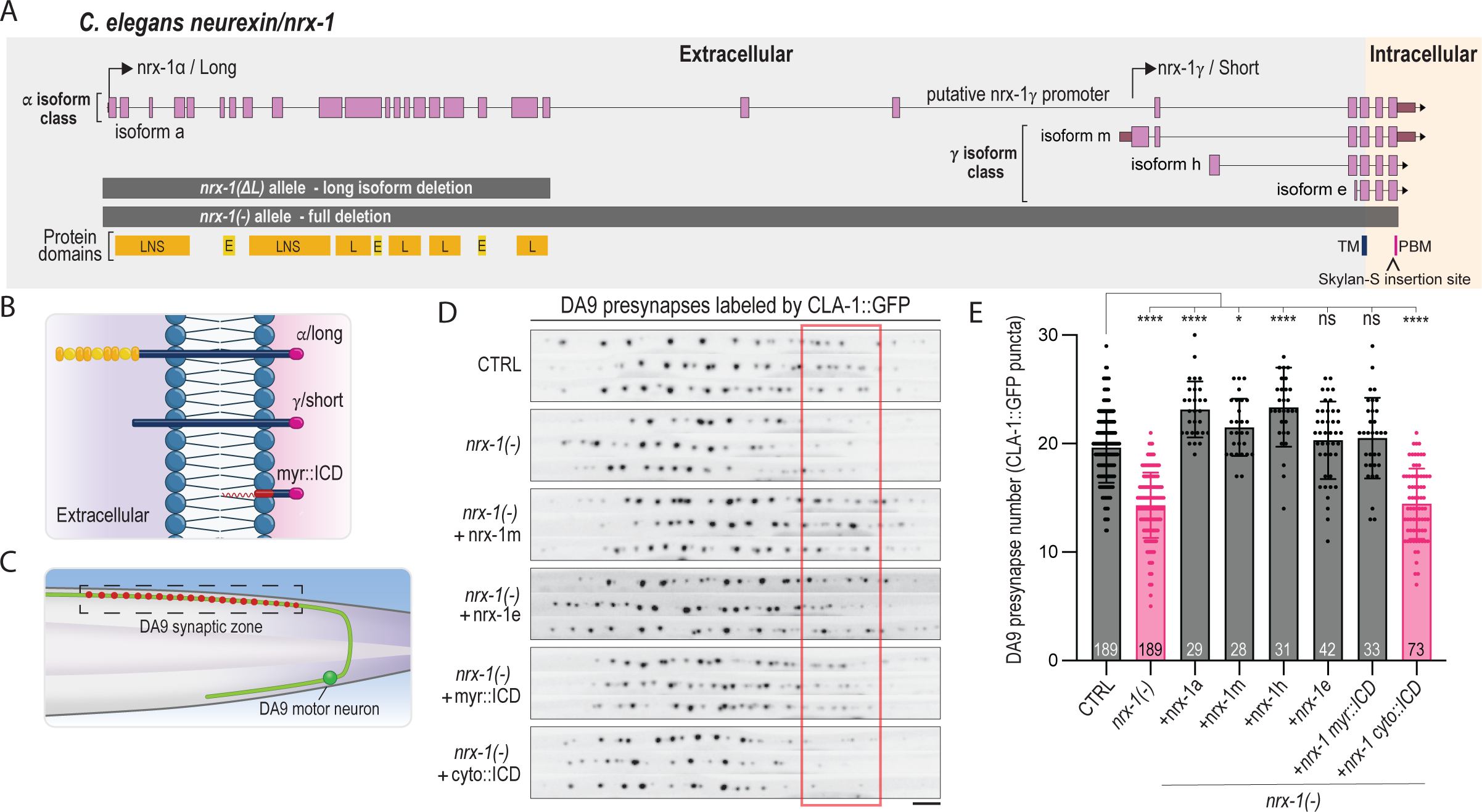
Synapse stabilization functions of NRX-1 are mediated by its intracellular domain. 1A. Schematic of *C. elegans neurexin/nrx-1* gene, isoforms, mutant alleles, and protein domains. The *nrx-1* gene encodes both α/long isoforms and γ/short isoforms. Protein regions encoded by the *nrx-1* gene are annotated at the bottom of the schematic (L/LNS = laminin-α, neurexin, sex hormone-binding globulin domain; E/EGF = EGF-like repeats; TM = transmembrane; PBM = PDZ binding motif). The fluorescent protein Skylan-S was inserted immediately prior to the C-terminal PBM. 1B. NRX-1 protein isoform schematics. The α/long isoform of NRX-1 has an extracellular domain with protein interaction motifs (yellows). The γ/short isoform protein has a small extracellular domain with no canonical protein interaction domains. The myr::ICD allele was engineered by removing extracellular sequence and replacing the transmembrane domain of *nrx-1* with a myristoylation sequence (red). 1C. Schematic of the DA9 excitatory motor neuron. The axon of DA9 forms a stereotypical set of synapses (red) in a specific position (box). 1D. DA9 presynapses (labeled by CLA-1::GFP) are reduced in *nrx-1(-)* animals, with puncta chiefly lost from the posterior region of DA9’s synaptic zone (indicated by a red box). DA9-specific expression of *nrx-1* isoform transgenes were assessed for rescue of the *nrx-1(-)* loss of presynapse phenotype. Three representative maximum intensity projections of the DA9 synaptic zone are stacked together for each genotype. Scale, 5 μm. 1D. Quantification of DA9 presynaptic puncta number in rescues of *nrx-1(-).* Sample size (number of worms) is indicated at the base of each bar, and individual values from each worm are represented as scatter points over each bar. Bars represent mean synapse number ± SD. (*p< 0.05, **p<0.01, ****p<0.001, Brown-Forsythe and Welch ANOVA tests with Dunnett’s T3 multiple comparisons test).

There are several splice isoforms of NRX-1(γ) that differ in their extracellular domains but contain an invariant intracellular domain (Fig. 1A). To determine whether particular extracellular motifs were important for synapse stabilization, we tested the ability of the NRX-1(γ) splice isoforms with the most dissimilar extracellular domains to rescue the *nrx-1(-)* phenotype in DA9. We found that DA9 neuron-specific expression of nrx-1(m), nrx-1(h), and nrx-1(e) splice isoforms of NRX-1(γ), whose extracellular sequences each decrease in length, respectively, all result in the rescue of CLA-1 puncta number in the *nrx-1(-)* background (Fig. 1D,E). This includes isoforms lacking predicted glycosylation motifs that have been shown in vertebrates to be important for neurexin transsynaptic functions^31^. The fact that even the nrx-1(e) isoform, which has a very short extracellular sequence (19 amino acids), was still able to restore presynaptic puncta (Fig. 1D,E), suggested that the extracellular domain of NRX-1 is not required for synapse maintenance. To test this, we expressed only the intracellular domain of NRX-1, tethered to the plasma membrane with a myristoylation sequence in the DA9 neuron (myr::ICD; schematized in Fig. 1B). We found that expression of the intracellular domain of neurexin was also able to rescue the loss of presynapses in *nrx-1(-)* mutants (Fig. 1D,E). However, rescue was dependent on its membrane localization as a cytosolic version (cyto::ICD) did not rescue the *nrx-1(-)* phenotype (Fig. 1D,E).

Altogether, these data indicate that the intracellular domain of neurexin, localized to the plasma membrane, is sufficient to mediate the synapse-stabilization function of neurexin.

### Neurexin clustering at presynaptic active zones does not require extracellular interactions

The ability of NRX-1(γ) and NRX-1(myr::ICD) to rescue the *nrx-1(-)* phenotype (Fig. 1) suggested that NRX-1(γ) isoforms may be able to cluster and contribute to presynaptic stabilization in the absence of extracellular or trans-synaptic interactions. To test this hypothesis directly, we examined the localization of endogenously labeled NRX-1 in living animals (see Fig. 2A for a schematic of the *C. elegans* body layout and imaging regions). We used a previously generated allele containing the Skylan-S fluorescent protein inserted into the shared intracellular domain of both isoforms, just prior to the PDZ-binding domain, in the endogenous *nrx-1* locus^16^ (annotated in Fig. 1A). In this allele, which we refer to as CTRL/*nrx-1::Skylan*, both NRX-1(γ) and NRX-1(α) isoforms are labeled, and we detect an average of ∼90 neurexin puncta per 100 μm in the axons of multiple neurons along the dorsal nerve cord (Fig. 2B,C; see Supp. Fig. 1B for examples of puncta detection). Seventy percent of the NRX-1 puncta area overlaps with endogenously-tagged active zone marker CLA-1::mScarlet (Supp. Fig. 2A). Loss of the sole homolog of neurexin’s canonical trans-synaptic binding partner neuroligin did not reduce endogenous NRX-1::Skylan clustering (Supp. Fig. 2B,C), suggesting that neurexin-neuroligin interactions are not required to cluster neurexin at synapses, and consistent with our previous observation that presynaptic assembly occurs normally in *C. elegans nlg-1(-)* mutants^16^.

**Figure 2.**
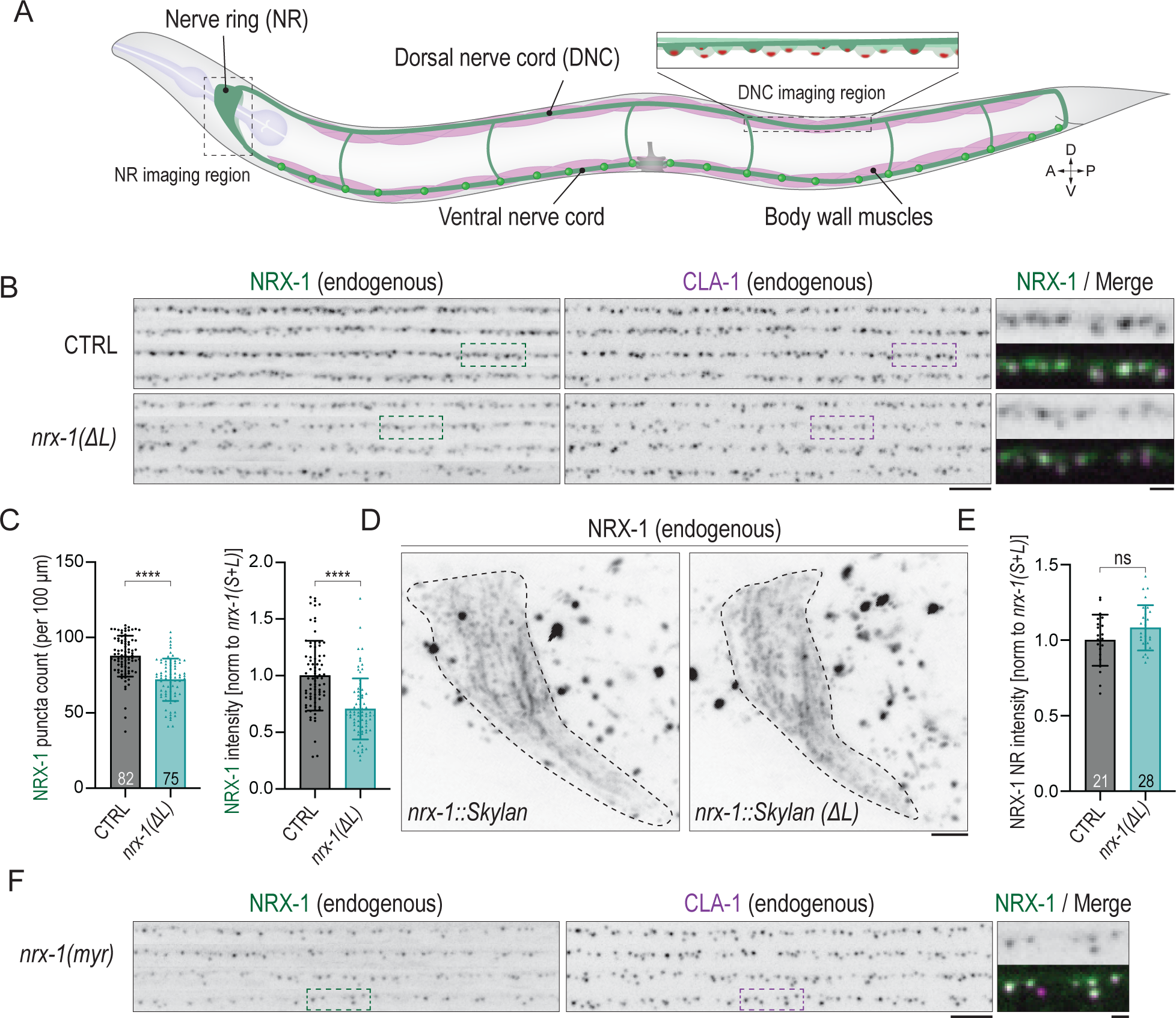
NRX-1 synaptic localization is mediated by its intracellular domain. 2A. Schematic of the *C. elegans* body plan, with imaging regions used in this paper indicated by dashed boxes. 2B. Endogenously labeled NRX-1::Skylan and CLA-1::mScarlet in CTRL and *nrx-1(ΔL)* animals. Merged images show magnifications of dashed box regions with NRX-1::Skylan pseudocolored in green and CLA-1::mScarlet in magenta. Scale, 5 μm. Merge scale, 1 μm. 2C. Quantification of NRX-1::Skylan puncta number and intensity in neurexin isoform alleles. 2D. Representative maximum projection images of endogenously labeled NRX-1 isoforms in the nerve ring in the heads of L4-stage animals. The dotted line indicates the boundary of the nerve ring. Saturated spots outside the nerve ring are autofluorescent granules. Scale, 5 μm. 2E. Quantification of mean NRX-1::Skylan isoform intensity in the nerve ring. 2F. Representative images of a strain expressing endogenous CLA-1::mScarlet and endogenous myristoylated intracellular domain of NRX-1::Skylan. Scale, 5 μm. Merge scale, 1 μm.

To visualize NRX-1(γ) alone, we deleted the genomic region encoding the long isoform in the endogenously tagged *nrx-1::Skylan* background to generate the *nrx-1::Skylan(ΔL)* allele (Fig. 1A). We observed that the remaining endogenously tagged NRX-1(γ) isoform (in the *nrx-1::Skylan(ΔL)* allele) still clusters at CLA-1-labeled presynaptic active zones in the dorsal nerve cord (Fig. 2B,C), indicating that it does not depend on transsynaptic interactions mediated by the long isoform in order to cluster. Surprisingly, the number of remaining NRX-1(γ)-only puncta (an average of 72 clusters per 100 μm) and the amount of protein at those clusters (as indicated by fluorescence intensity) was only mildly reduced relative to the *nrx-1::Skylan* allele containing both isoforms (Fig. 2B,C), suggesting that the short isoform is the dominant isoform at dorsal nerve cord synapses. To determine whether the short isoform also predominates at synapses throughout the nervous system, we compared neurexin fluorescence intensity in the nerve ring neuropil of *nrx-1::Skylan* and *nrx-1::Skylan(ΔL)* animals and observed no significant difference, suggesting that the short isoform is the predominantly expressed isoform at synapses throughout the *C. elegans* nervous system, or alternatively can be upregulated to compensate for loss of the long isoform (Fig. 2D,E).

To test whether the remaining extracellular or transmembrane domains of NRX-1(γ) are required for its clustering, we deleted these domains from the *nrx-1::Skylan(ΔL)* allele and replaced them with an N-terminal myristoylation tag (generating the *nrx-1::Skylan(myrICD)* allele). Though reduced in intensity, we found that the myristoylated short isoform, NRX-1(myr), still clusters and localizes to CLA-1-labeled active zones (Fig. 2F), providing strong evidence that neurexin localization and clustering do not depend on extracellular interactions.

### Intracellular PDZ-based interactions cluster neurexin at presynaptic sites

In light of our results indicating that neurexin does not require extracellular interactions to localize to presynaptic sites or to stabilize synapses, we undertook a candidate screen of presynaptic active zone genes to identify regulators of neurexin localization. Most genes had little to no effect on NRX-1 localization or clustering (Supp. Fig. 3A-D)^28,33^. We found that *unc-104/kinesin-3* mutants exhibited a severe loss of neurexin protein within the axons of the dorsal nerve cord, consistent with previous work showing that this molecular motor is responsible for NRX-1 trafficking^32^. However, the remaining neurexin present in the dorsal nerve cord of *unc-104(-)* mutants retained its ability to form clusters (Supp. Fig. 3E,F). In contrast, we found that the loss of the active zone scaffold gene *syd-1* caused a severe defect in neurexin clustering (Fig. 3A-C, Supp. Fig. 3A,B). We also found more subtle phenotypes in mutants of the active zone genes *syd-2/Liprin-α* and *nab-1/Neurabin* (Supp. Fig. 3A,B), known regulators of SYD-1 function^18,33^.

**Figure 3.**
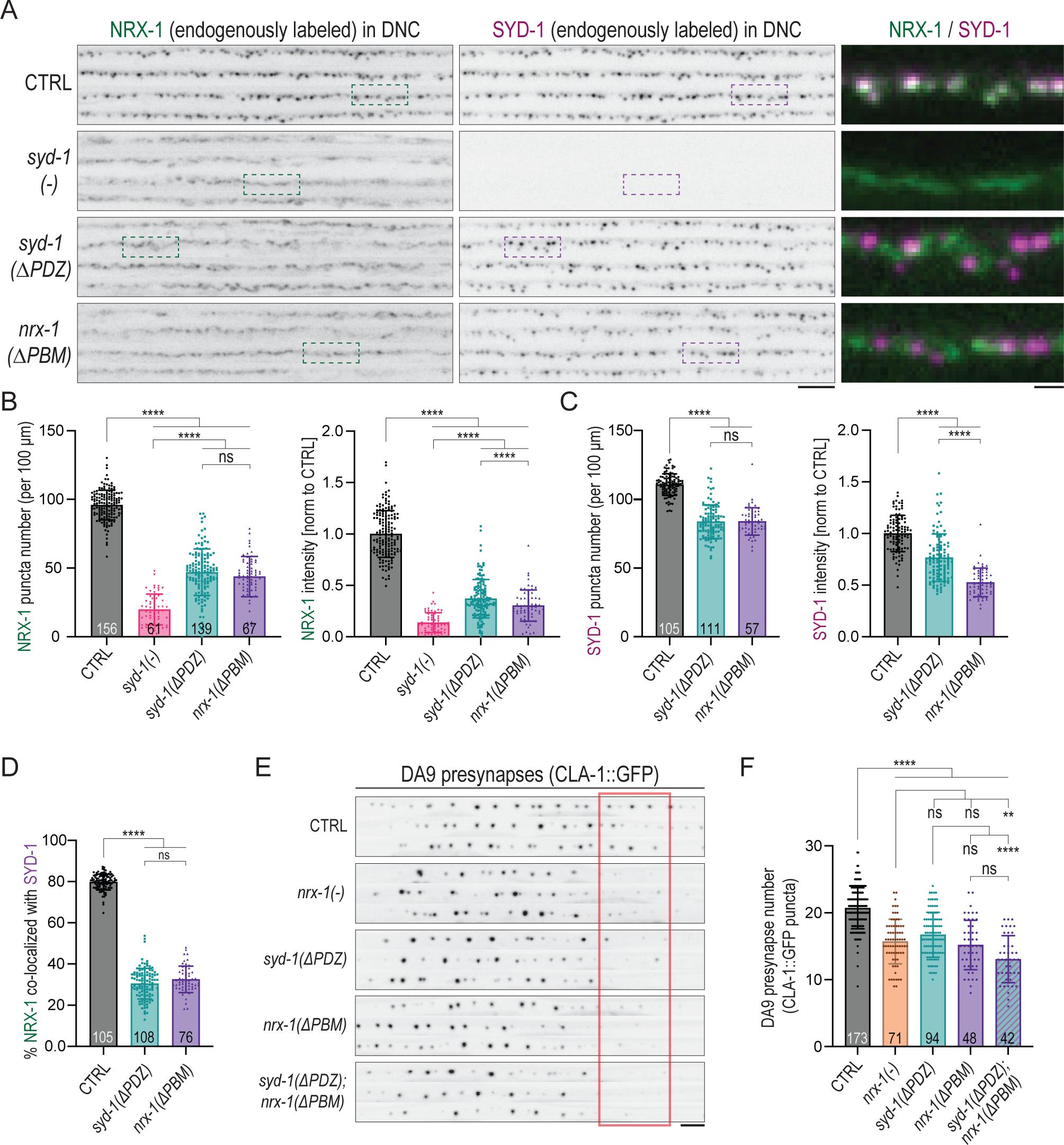
Intracellular scaffold protein SYD-1 recruits neurexin to presynapses using a PDZ interaction. 3A. Representative images of endogenously tagged NRX-1::Skylan and SYD-1::mScarlet in mutants that disrupt the PDZ-based interaction. Note that NRX-1 declusters in all three mutant genotypes, while SYD-1 remains competent to cluster in both the *nrx-1(ΔPBM)* and *syd-1(ΔPDZ)* backgrounds. The *syd-1(-)* allele does not contain CDs to express a fluorescent protein. Scale, 5 μm. Merge scale, 1 μm. 3B,C. Quantification of NRX-1::Skylan (B) or SYD-1::mScarlet (C) puncta number and intensity in control and mutant alleles of *syd-1* and *nrx-1*. 3D. Quantification of % NRX-1 ROI area co-localized with SYD-1 in *syd-1(ΔPDZ)* and *nrx-1(ΔPBM)*. 3E. CLA-1::GFP-labeled presynapses in the DA9 neuron of *nrx-1(-), nrx-1(ΔPBM), syd-1(ΔPDZ),* and *nrx-1(ΔPBM); syd-1(ΔPDZ)* mutants. Scale, 5 μm. 3F. Quantification of DA9 presynapse number in mutants that disrupt the PDZ interaction between NRX-1 and SYD-1.

NRX-1::Skylan clusters colocalized tightly with endogenously tagged SYD-1::mScarlet (Fig. 3A,D). To ablate potential interactions between NRX-1 and SYD-1^29^, we removed the PDZ domain of SYD-1 from the endogenous locus of the *syd-1::mScarlet* allele and assessed NRX-1 and SYD-1 localization. The resulting *syd-1(ΔPDZ)* mutation caused a dispersion of NRX-1 (Fig. 3A,B) and loss of colocalization between NRX-1 and SYD-1 (Fig. 3A,D). SYD-1(ΔPDZ) itself remained clustered, albeit at slightly reduced levels (Fig. 3A,C). In a complementary experiment, the removal of the neurexin C-terminal PDZ domain-binding motif (PBM) to generate the *nrx-1::Skylan(ΔPBM)* allele recapitulated the NRX-1 dispersion phenotype (Fig. 3A,B,D). In this allele, SYD-1 puncta number and intensity were also reduced (Fig. 3A,C), indicating that while the SYD-1 PDZ domain is absolutely critical for NRX-1 clustering, NRX-1 also plays a role in stabilizing SYD-1 clusters.

To determine the contribution of the PDZ-based interaction between NRX-1 and SYD-1 to synapse stability, we evaluated the number of CLA-1-labeled active zones in the DA9 neuron and found that *syd-1(ΔPDZ)* and *nrx-1(ΔPBM)* mutants were indistinguishable from that of *nrx-1(-)* mutants (Fig. 3E,F). Double mutants of *syd-1(ΔPDZ)* and *nrx-1(ΔPBM)* were not significantly more severe than the *nrx-1(ΔPBM)* single mutant (Figure 3E,F). These data suggest that SYD-1 and NRX-1 function together in the same pathway, through a PDZ-based interaction, to stabilize presynapses.

### SYD-1 and SYD-2 clustering at nascent synapses precedes that of NRX-1, but NRX-1 is required for subsequent synapse stabilization

Our finding that NRX-1 clustering is dependent on SYD-1, but SYD-1 can cluster in the absence of NRX-1, suggested that SYD-1 might arrive at nascent presynaptic specializations prior to neurexin and facilitate its recruitment to those sites. To test this prediction and determine the order in which these proteins arrive at presynapses, we imaged the developing embryonic nerve ring in a strain expressing both endogenously-tagged NRX-1::Skylan and SYD-1::mScarlet. Imaging in the embryonic nerve ring enabled us to observe the formation of the earliest synapses being established. In separate animals, we also imaged endogenously tagged SYD-2/Liprin-α, which is detectable in the nerve ring very early in *C. elegans* embryonic development (bean stage, ∼360 minutes post-fertilization, mpf) and is thought to contribute to presynaptic assembly by forming phase-separated condensates that can recruit additional presynaptic proteins^23^. We found that SYD-1::mScarlet first becomes detectable in the nerve ring at the 2-fold embryonic stage (∼490 mpf), at which point, in the same animal, NRX-1::Skylan remains undetectable (Fig. 4A). NRX-1::Skylan first becomes visible in the nerve ring at the 3-fold stage (∼550 mpf) of embryonic development (Fig. 4A), the same stage at which synaptic vesicle markers such as RAB-3 have previously been shown to appear^23^. To confirm that this result was not due to differences in the brightness or maturation time of the respective fluorophores, we validated this result in singly-labeled NRX-1::GFP and SYD-1::GFP strains (Supp. Fig. 4A).

**Figure 4.**
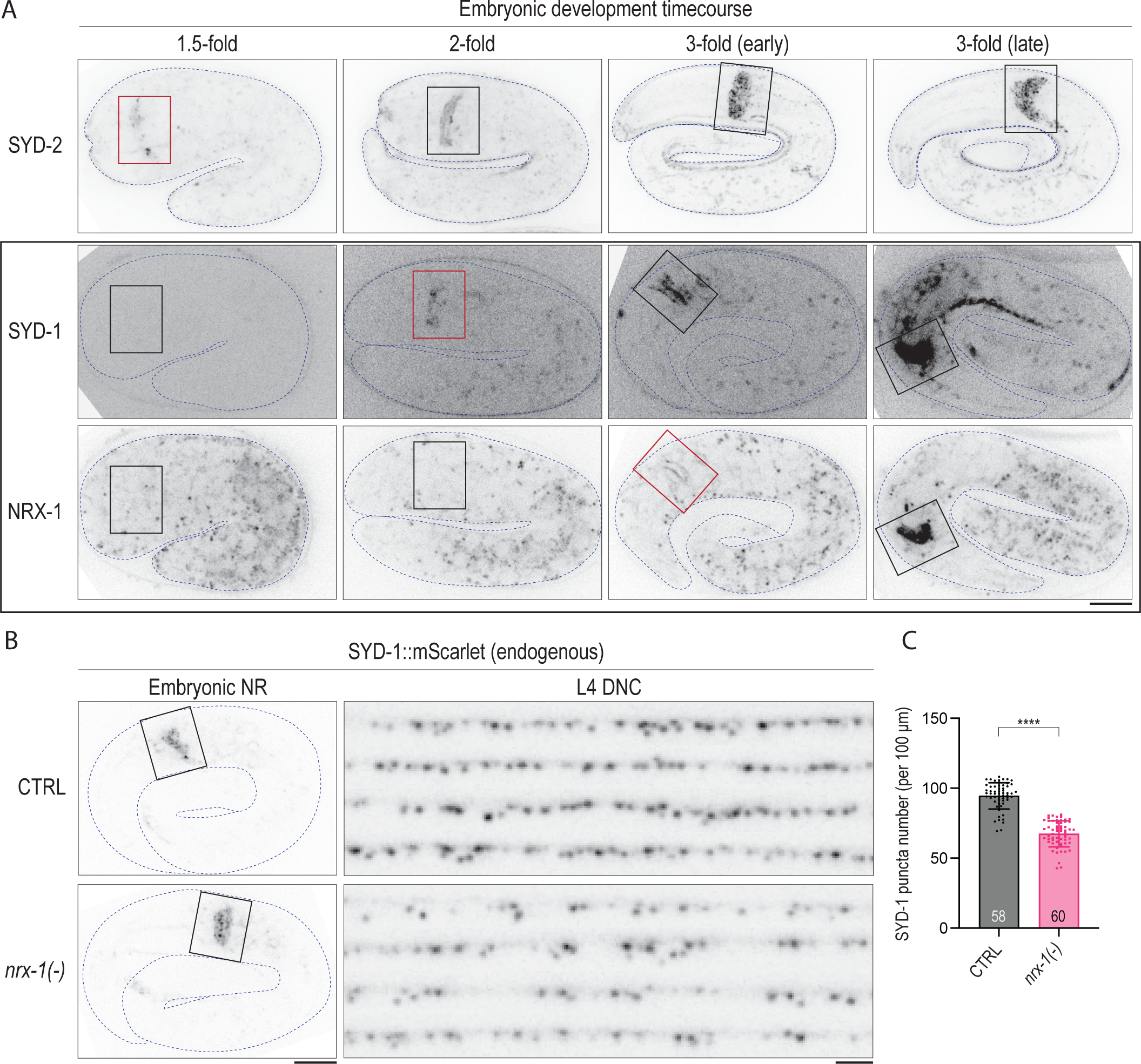
SYD-1 accumulation precedes that of NRX-1 at nascent presynapses. 4A. Embryos at different developmental timepoints expressing endogenously labeled SYD-2, SYD-1, and NRX-1. The developing nerve ring at each stage is enclosed with a box (colored red at the stages in which each protein is first detected). SYD-1 and NRX-1 were imaged together in the same animals, and single planes of interest from confocal z-stacks are displayed. Scale, 10 μm. 4B,C. Endogenously labeled SYD-1 in the embryonic nerve ring (boxed), left, and in the dorsal nerve cord of L4-stage animals (right). Embryo scale, 10 μm. L4 DNC scale, 2.5 μm. 4C. Quantification of SYD-1 puncta in a *nrx-1* mutant in the L4-stage DNC.

SYD-1 accumulation at synapses in the embryonic nerve ring is not impacted by the loss of *nrx-1* (Fig. 4B), in concordance with the observation that SYD-1 accumulation occurs prior to the arrival of NRX-1 (Fig. 4A). However, by the last larval stage of development (L4), SYD-1 puncta number are decreased in the dorsal nerve cord of *nrx-1* mutants (Fig. 4B,C), suggesting that while NRX-1 is not required for the initial expression or localization of SYD-1 to presynapses, it is necessary for SYD-1’s long-term stability at those sites. Loss of *nrx-1* had a similar effect on SYD-2, with no impact on SYD-2::GFP accumulation in the embryonic nerve ring but a significant loss of puncta number and intensity in L4 animals (Supp. Fig. 4B,C), in agreement with prior results^16^. Mutants disrupting the liquid-liquid phase separation properties of SYD-2/Liprin-α or ELKS-1^23^ had no distinguishable impact on NRX-1 puncta number or intensity in the dorsal nerve cord of L4 animals (Supp. Fig. 4D,E), consistent with the previous finding that SYD-2/Liprin-α phase separation has no effect on the presynaptic localization of SYD-1^23^.

Together, these data contradict canonical models in which cell-adhesion molecules recruit cytosolic scaffolds to presynapses. Instead, our results show that the cytosolic scaffold molecule SYD-1 clusters at nascent presynaptic specializations prior to and independently of NRX-1, and then recruits NRX-1. NRX-1 is subsequently required for maintaining and stabilizing those active zone clusters.

### Computational modeling predicts SYD-1-membrane interaction via its C2 and PDZ domains

We found that SYD-1 precedes NRX-1 to nascent presynapses and is responsible for neurexin’s clustering at these sites, but how SYD-1 localizes to the plasma membrane to cluster NRX-1 is unknown. SYD-1 contains a C2 domain, a known membrane-targeting module present in many proteins^34–37^. To determine if the C2 domain of SYD-1 might be capable of targeting SYD-1 to the plasma membrane in the absence of a transmembrane binding partner, we performed both course-grained and atomistic molecular dynamics (MD) simulations to computationally predict whether the SYD-1 C2 domain can interact with the cell membrane. We predicted the SYD-1 C2 domain structure with AlphaFold and initiated 20 independent trajectories with the C2 domain rotated at a random angle relative to an anionic membrane of known composition (75% POPC, 15% POPS, 5% PIP_2_, 5% cholesterol) previously used in MD simulations^38^. In all simulations, the SYD-1 C2 domain stably bound to the membrane within 100 ns (Fig. 5A,B), mediated by interactions between six basic amino acids within the C2 domain and the phosphates of PIP_2_ (Supp. Fig. 5A). This binding did not occur in simulations with membrane lacking PIP_2_ (Fig. 5C). We used DDmut^39^ to predict the change in Gibbs free energy (ΔΔG) induced by mutation of all six of the PIP_2_-interacting residues to alanines and found that these changes are not predicted to be disruptive to the stability of the C2 domain structure (Supp. Fig. 5B).

**Figure 5.**
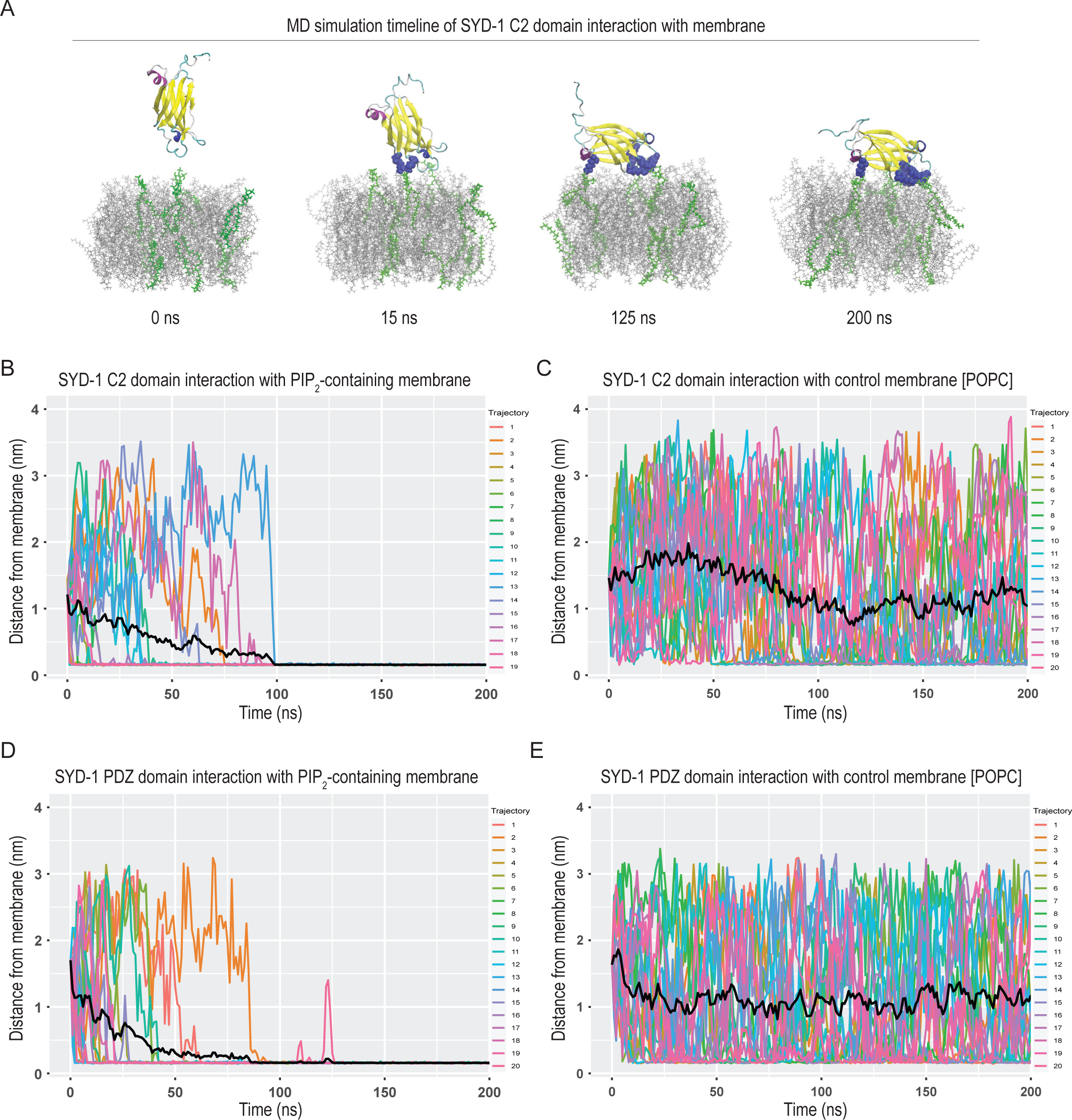
MD simulations predict interactions between SYD-1 C2 and PDZ domains and PIP_2_ phospholipids. 5A. Snapshots from an atomistic MD simulation in which an Alphafold-predicted SYD-1 C2 domain structure was tested for interaction with a theoretical membrane over the course of 200 nanoseconds. The amino acids participating in interactions with the membrane are indicated in blue, and the PIP_2_ phospholipids in the membrane are green. 5B-E. The minimum protein-lipid distance as a function of time is represented by a different color for each of the trajectory repeats per simulation condition, with black lines representing a rolling average distance between the domain and membrane at each time point.

Based on reports that PDZ domains are also able to bind to membrane lipids (for example, in Syntenin^40–42^, Par3^43^, and PICK1^44,45^), we modeled the SYD-1 PDZ domain in MD simulations. We found that the SYD-1 PDZ domain was also predicted to stably bind to membrane within about 125 nanoseconds in all 20 simulations, again in a manner dependent on the presence of PIP_2_ (Fig. 5D,E). In contrast, MD simulations using the same conditions did not predict any stable interaction between the SYD-1 RhoGAP domain and membrane within 200 nanoseconds (Supp. Fig. 5C).

Next, we modeled the structure of the interaction between the SYD-1 PDZ domain and NRX-1 PBM at the plasma membrane (represented in Supp. Fig. 5D). We applied a coarse-grained Langevin dynamics (CGLD) simulation method (represented in Supp. Fig. 5E) to predict how this interaction can be affected by the attachment of the SYD-1 PDZ and C2 domains to the membrane. The simulation results showed that the interactions between the SYD-1 PDZ domain and NRX-1 PBM are generally increased as the probability of attachment of the SYD-1 PDZ and C2 domains to the membrane are increased (Supp. Fig. 5F), suggesting that initial membrane interactions by SYD-1 may subsequently lead to more stable protein-protein interactions between SYD-1 and NRX-1. Together these data suggest a mechanism by which SYD-1 may initially interact with the plasma membrane through its C2 and/or PDZ domains, and thus position it to cluster the transmembrane protein NRX-1.

### SYD-1 localization *in vivo* is dependent on both its C2 and PDZ domains

To test our modeling predictions and determine whether the PDZ and C2 domains of SYD-1 contribute to its stability at presynaptic sites *in vivo*, we removed each of these domains individually from the endogenous *syd-1::mScarlet* allele. The *syd-1(ΔC2)* mutation led to a mild but significant reduction in the puncta number and intensity of both SYD-1 and NRX-1, and a minor reduction in the co-localization between SYD-1 and NRX-1 (Fig. 6A-D). As in Fig. 3, *syd-1(ΔPDZ)* led to a small decrease in the number of SYD-1 puncta and a reduction in the intensity of SYD-1 at remaining sites, as well as a severe declustering of NRX-1 and loss of co-localization (Fig. 6A-D). Because the single domain deletions of *syd-1(ΔPDZ)* or *syd-1(ΔC2)* do not drastically reduce SYD-1 clustering, deletion of each of these domains does not appear to have a major effect on the folding, stability, or trafficking of SYD-1. Instead, these two domains appear to tune SYD-1’s clustering behavior: The C2 domain is required for normal puncta size/intensity, and the PDZ domain for maintaining appropriate puncta number (consistent with the role of its PDZ binding partner, NRX-1, in maintaining the correct number of presynapses). Removal of both the SYD-1 C2 and PDZ domains led to a severe reduction in SYD-1 puncta number and intensity (Fig. 6A,C), indicating that these two domains together are largely responsible for SYD-1 localization and clustering. Similarly, while loss of either the SYD-1 C2 or PDZ domains had minor effects on CLA-1::GFP-labeled presynapses in DA9, loss of both domains together led to a severe reduction in presynaptic puncta that was not significantly different from the *syd-1(-)* allele (Supp. Fig. 6A,B). Moreover, deletion of the SYD-1 PDZ domain in combination with mutations of the six predicted PIP_2_-interaction residues of the C2 domain (C2*; Supp. Fig. 5A) led to a reduction In SYD-1 puncta number and intensity that was almost as severe as deleting the entire C2 domain (Fig. 6A-D), suggesting that the effect of the double domain deletion was largely due to the removal of those six residues. The severe reduction of SYD-1 intensity is not due to the loss of presynaptic clusters as a whole, as SYD-2/Liprin-α still remains clustered (Supp. Fig. 6C,D). However, SYD-2 is reduced in both *syd-1(ΔPDZ ΔC2)* and *syd-1(-)* mutants to similar extents (Supp. Fig. 6C,D).

**Figure 6.**
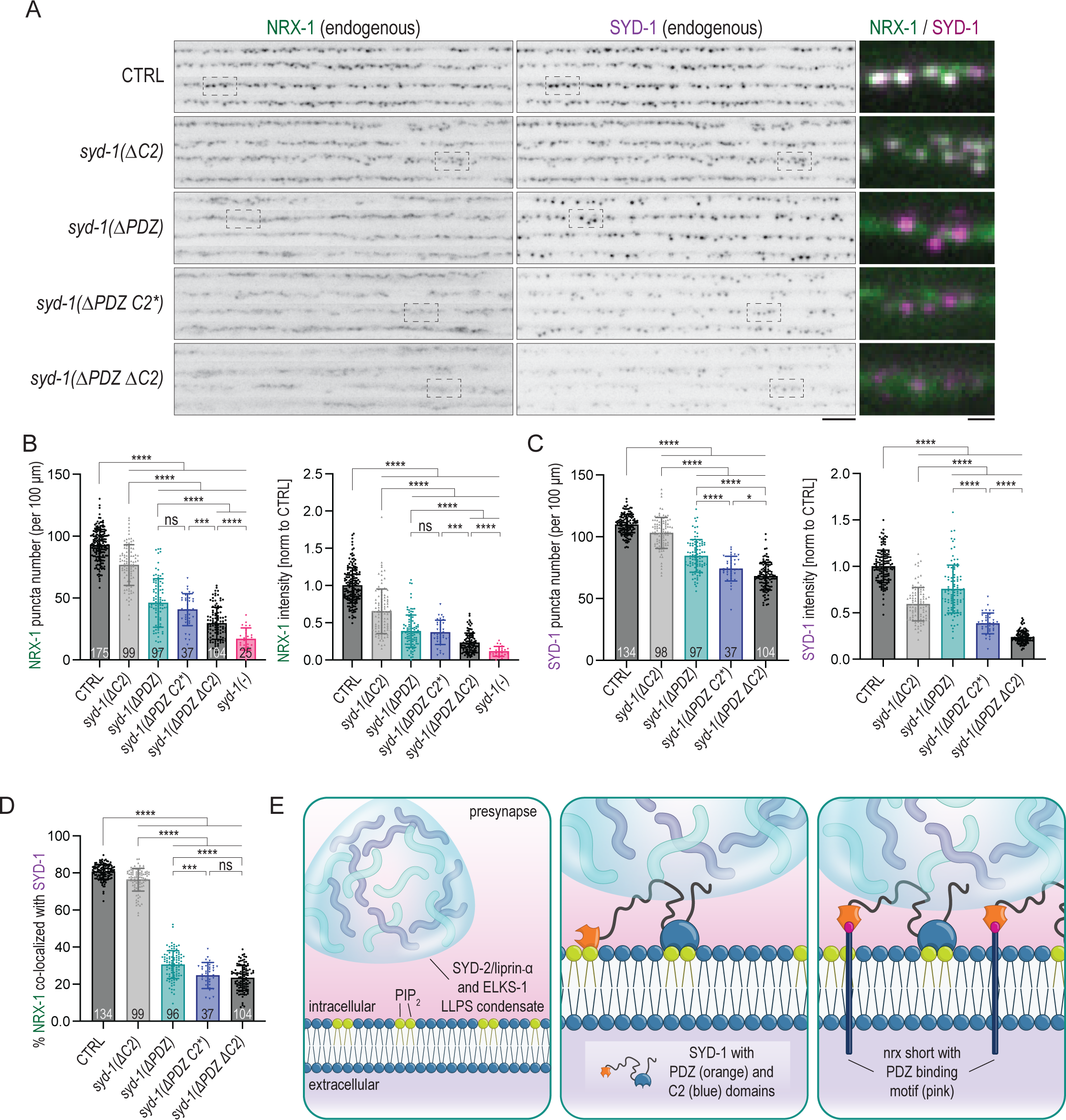
The C2 and PDZ domains of SYD-1 redundantly mediate SYD-1 synaptic localization *in vivo*. 6A. Representative images of endogenously labeled SYD-1 lacking its C2 or PDZ domains (or both), as well as the *syd-1(ΔPDZ C2*)* allele, in which the PDZ domain was removed and the predicted PIP_2_-interacting residues of the C2 domain were mutated to alanine. Dashed boxes indicate regions used for merged magnifications at right, which depict NRX-1 in green and SYD-1 in magenta. Scale, 5 μm. Merge scale, 1 μm. 6B. Quantifications of NRX-1 puncta number and puncta intensity in *syd-1* mutant alleles. 6C. Quantification of SYD-1 puncta number and puncta intensity in *syd-1* mutant alleles. 6D. Quantification of co-localization between endogenous NRX-1::Skylan and SYD-1::mScarlet in *syd-1* mutant alleles. 6E. The presynaptic scaffold protein SYD-1 coordinates synapse assembly through interactions with PIP_2_ and subsequently recruits downstream synapse components, including neurexin. Early in synapse development, SYD-2/Liprin-α and ELKS-1 form phase-separated condensates that recruit and concentrate downstream active zone proteins, enabling them to participate in the multivalent interactions necessary to build and organize functional presynapses (panel 1). Following the expression of SYD-2 and ELKS (but prior to most other active zone proteins), SYD-1 protein becomes detectable at nascent presynapses. SYD-1 is recruited to the plasma membrane through interactions between its C2 and PDZ domains and PIP_2_ phospholipids (panel 2). SYD-1 may recruit the phase-separated condensate through interactions with SYD-2 and ELKS-1 (panel 2), and coordinates the recruitment of NRX-1/neurexin to presynapses through PDZ-based interactions (panel 3). The γ /short isoform of NRX-1 is the isoform predominantly expressed by the *C. elegans* nervous system and functions in synapse stabilization.

Taken together, these *in vivo* data confirm the *in silico* predictions of our computational modeling by demonstrating that the C2 domain and the PDZ domain of SYD-1 have redundant roles in SYD-1 localization. Moreover, we identify interactions with membrane PIP_2_ as a potential mechanism for SYD-1 localization to the presynaptic membrane in the absence of transmembrane cell-adhesion molecules like NRX-1.

## DISCUSSION

Our current understanding of the contributions of synaptic CAMs like neurexin to synaptogenesis remains rudimentary and rooted in the idea that trans-synaptic interactions serve to initiate this process. However, our results challenge this paradigm. We show that the γ/short neurexin isoform can cluster *in vivo* in the absence of extracellular interactions and that the intracellular domain of neurexin, tethered to the membrane, is sufficient to perform synapse stabilization functions. Moreover, we determined that the intracellular scaffold protein SYD-1 acts as the principal regulator of neurexin’s synaptic localization through a PDZ-based interaction. While SYD-1 and its PDZ domain are critical to neurexin clustering, SYD-1 does not depend on neurexin nor its own PDZ domain to cluster at presynapses. These results suggest that SYD-1 is positioned upstream of neurexin in a synapse assembly pathway. In support of this concept, we found that the initial accumulation of SYD-1 at nascent presynapses precedes the arrival of neurexin. Finally, we investigate the mechanism by which SYD-1 localizes and clusters independently of neurexin. We find that SYD-1 recruitment to synapses is dependent on its C2 and PDZ domains, both of which we computationally predict to interact with the phospholipid PIP_2_ in the plasma membrane. Taken together, our results reveal a novel mechanism of presynaptic development in which cytosolic scaffold proteins like SYD-1 serve to initiate the formation of presynaptic assemblies at the plasma membrane by first interacting with membrane phospholipids and subsequently recruiting transmembrane molecules such as neurexin to presynaptic sites (Fig. 6E). Crucially, we identify a pivotal role of the evolutionarily conserved yet largely-uncharacterized non-trans-synaptic γ/short isoform of neurexin in stabilizing these nascent presynaptic assemblies at the membrane.

### The role of the neurexin intracellular domain in synapse stabilization

Neurexins are a class of synaptic CAMs that have been implicated in schizophrenia, autism, and intellectual disability^46–48^, and thus represent clinically relevant treatment targets whose functions in synapse development pathways have not yet been firmly established. The canonical role of CAMs—including neurexin—is in the initiation of synapse assembly (based on *in vitro* synapse formation assays in cell culture^49–51^). However, our *in vivo* results contradict this model for neurexin function. We determined that endogenous neurexin clustering at presynapses is dependent on the intracellular protein SYD-1, and we detected SYD-1 before NRX-1 at early synapses, challenging the concept that CAMs interact trans-synaptically to initiate presynaptic assembly. We found that both the SYD-1 PDZ domain and the NRX-1 PDZ domain binding motif are critical for neurexin clustering at presynapses, demonstrating an intracellular mechanism for NRX-1 localization. Interestingly, large-scale *in vitro* screens to identify PDZ interactions in *C. elegans* identified SYD-1 as a likely partner to NRX-1^52^. As we observed in *C. elegans,* a PDZ-based interaction underlies binding between *Drosophila* neurexin and SYD-1 and is thought to aid in the retention of neurexin at presynapses^29^. However, in contrast to vertebrates, *Drosophila* lacks the γ/short isoform of neurexin, suggesting that worms may provide a more suitable model for characterizing the function of the short, non-trans-synaptic γ isoform and the specific role it plays in presynaptic assembly in mammals.

Biochemical characterizations of mammalian neurexin have demonstrated that the neurexin intracellular domain directly interacts with CASK^53,54^, Mint^55^, FERM-domain protein 4.1^56^, syntenin^57^, and synaptotagmin^58^, providing potential links between neurexin and the pathways that regulate presynaptic organization, cytoskeletal assembly, Ca^2+^ channel clustering, synaptic vesicles, and exocytosis. Most of these interactions are mediated by neurexin’s C-terminal PBM and the PDZ domains of these proteins. Whether these intracellular interactors are necessary for neurexin localization and function *in vivo* is unknown. Research in the invertebrate *D. melanogaster* has identified other PDZ domain-mediated interactions between neurexin’s intracellular domain and presynaptic intracellular proteins, including active zone protein Spn^59^ (NAB-1 in *C. elegans)*. In line with canonical models of synapse assembly, these proteins have been assumed to be recruited by neurexin, although this has not been demonstrated.

Our results highlight the critical role for neurexins in synapse maintenance rather than in synapse initiation. We have previously shown that the γ/short isoform of neurexin, which is present but largely uncharacterized in mammals, is sufficient to stabilize presynapses and prevent them from being eliminated by inhibitory cues in *C. elegans*^16^. Here we find that endogenous NRX-1 γ/short can still cluster upon removal of its short extracellular sequence, and that the neurexin intracellular domain, when tethered to the plasma membrane, is sufficient to stabilize presynapses against elimination. We propose that synapse stabilization, which is mediated by the intracellular domain common to all neurexin isoforms, represents the core, conserved function of neurexin. A fundamental question for future exploration is how neurexins mediate synapse stabilization, which they may accomplish through regulation of the actin cytoskeleton^56,60,61^.

Our efforts to identify the role of the γ/short isoform of neurexin led us to determine that it is likely the more abundantly expressed isoform in the nervous system of *C. elegans* at the late larval L4 stage. The wide expression of the γ/short isoform at this stage may signify a cost-saving measure by the nervous system to cheaply produce a protein that accomplishes the base function of synapse stabilization. While the α/long isoform of neurexin contains the same intracellular domain and is thus able to mediate synapse stabilization, its expression may instead be reserved for additional specialized roles that require extracellular input in particular developmental or regulatory contexts. For example, the α/long isoforms of neurexin have been implicated in neurite outgrowth/axonal guidance^62–64^ and postsynaptic clustering of AChRs^65^. How the expression of the α/long and γ /short isoforms is differentially regulated in the nervous system through development and what guides the logic of neurexin isoform usage represent interesting questions that can be addressed in the simple *C. elegans* system.

### An intracellular mechanism for presynaptic assembly

Despite mounting *in vivo* evidence from mammalian knock-out models that individual CAMs are responsible for functional maturation of synapses rather than their initiation^4,11,66,67^, redundancy within the large class of CAMs is often invoked to explain this apparent discrepancy between model and data. However, evidence for completely cell-intrinsic mechanisms of pre- and postsynaptic assembly can be found in the literature. The vertebrate neuromuscular junction arranges neurotransmitter receptors into pre-patterned clusters long before the arrival of the motor neuron^68^. Cultured mammalian neurons can release synaptic vesicles from nascent active zones before reaching any target cell^69^. *Drosophila* mutants in which muscles fail to develop still exhibit ultrastructurally normal presynaptic specializations without any opposing postsynaptic structure^70^. Together, this evidence implies that transsynaptic interactions in general may not be required for the initial clustering of pre- and postsynaptic components. The lack of an alternative model for synapse assembly has hindered the acceptance of this conclusion. Our results identify a mechanism by which intracellular proteins can cluster at membranes in the absence of transmembrane proteins, and how these cell-intrinsic mechanisms can then subsequently be coupled to transsynaptic signaling.

SYD-1 and SYD-2 have long been known to play a central role in synapse assembly in invertebrates. In vertebrates, the intrinsically disordered region of mouse mSYD1a has been shown to interact with Liprin-α2 and munc18-1 to facilitate synaptic vesicle docking at the active zone^27^. Human SYD-1 homologs SYDE1 and SYDE2 have been implicated in intellectual disability and autism^71,72^ and have been predicted to function as signaling hubs for neuronal development^73^. Aside from the SYDE homologs, other mammalian proteins may perform similar functions as SYD-1 in *C. elegans*. For example, mammalian RIM, which contains a C2b domain that has been shown to target it to PIP_2_-containing membrane^37^ and a PDZ domain that coordinates Ca^2+^ channel localization^74^, may function similarly to invertebrate SYD-1 in clustering synapse-stabilizing transmembrane molecules during synapse assembly. The role of SYD-2/Liprin-α in active zone assembly is highly conserved in mammals, where it has been shown to recruit several components of the active zone release machinery and facilitate vesicle docking and neurotransmitter release^24–26^. Recent evidence has linked the gene PPFIA3, which encodes Liprin-α3 in humans, with developmental delay, intellectual disability, autism, and epilepsy^75^. Given their clinically relevant roles in regulating synapse assembly, characterizing how SYD-1 and SYD-2/Liprin-α—as well as their functional homologs—contribute to synaptogenesis is of critical importance.

In the molecular hierarchy of invertebrate synapse assembly, SYD-1 has been positioned upstream of SYD-2/Liprin-α, with reports that it promotes SYD-2’s localization and retention at active zones while not being affected by the loss of SYD-2^17–20,23,28,29,76^. SYD-1 and SYD-2 have been shown to interact in yeast two-hybrid assays^23^, and each also interacts with ELKS-1/Bruchpilot^19,23,28^, another scaffold protein that localizes to presynapses very early relative to other active zone proteins^19,23^. Recent evidence has implicated liquid-liquid phase separation of active zone proteins in presynaptic active zone assembly assembly^26,77,78^, including an *in vivo* demonstration of phase separation of SYD-2 and ELKS-1 at early stages of synapse development in *C. elegans*^23^. These condensates are thought to recruit and concentrate downstream active zone proteins, enabling them to participate in all the multivalent interactions necessary to build and organize the functional active zone. Although SYD-1 is positioned genetically upstream of SYD-2 in synapse assembly^18^, we detected SYD-2 in neurons of the developing embryonic nerve ring of *C. elegans* earlier than SYD-1. We propose a model, schematized in Figure 6E, in which synapse assembly occurs through the following sequential steps: (1) the early formation of the SYD-2/ELKS-1 phase-separated condensates, which incorporate other downstream active zone proteins as they are expressed, (2) the expression and localization of SYD-1 to both the SYD-2/ELKS-1 phase-separated condensate through its multivalent interactions with SYD-2 and ELKS and to PIP_2_-enriched regions of the axonal plasma membrane through its C2 and PDZ domains, and (3) SYD-1-mediated recruitment of the CAM neurexin through PDZ-based interactions. This model serves to explain both the temporal progression of the arrival of synaptic proteins, as well as the stabilization functions of both SYD-1 and neurexin on presynaptic assemblies later in development.

PIP_2_ is predominantly localized at the plasma membrane, is enriched in small raft-like structures^79–83^, and has been hypothesized to be enriched at presynaptic release sites^82,84^, where it is necessary for efficient vesicle fusion^85–87^. Our findings highlight the critical importance of protein-membrane interactions between SYD-1 and PIP_2_ in the recruitment of SYD-1 to nascent synapses, which enables SYD-1 to perform its functions in recruiting and organizing downstream synaptic proteins including both CAMs like neurexin and other presynaptic scaffolds like SYD-2/Liprin-α. Recent work in human iPSC-derived neurons has implicated the rare endosomal membrane phospholipid, PI(3,5)P_2_ in directing the transport of synaptic vesicle precursors to presynaptic domains^88^. In addition, PI(4,5)P_2_ was recently shown to regulate synaptic vesicle endocytosis downstream of synaptotagmin-1 at the plasma membrane^89^. Finally, cholesterol regulation was shown to be important broadly for regulating synapse number and function, through a process that may be dysregulated in Alzheimer’s disease^90^. Together these data indicate that membrane lipid composition is a critical, conserved, and perhaps overlooked mediator of synaptic development and function.

Our studies define a mechanism for the initiation of presynaptic assembly that positions a cytosolic scaffold protein interacting with the cell membrane upstream of a synaptic cell adhesion molecule. Together, this work highlights the importance of protein-lipid interactions in synapse formation, reframes the order in which proteins assemble at the synapse, and identifies a role for the γ/short isoform of neurexin in synaptic maintenance.

## ACKNOWLEDGEMENTS

We thank Hannes Bülow and members of the Bülow lab, Scott Emmons, David Hall, and members of the Kurshan lab for discussions during the course of this work and comments on the manuscript. Thanks also to Brian Mueller, Bryen Jordan, Kelsie Eichel and Callista Yee for helpful comments on the manuscript. We thank Kang Shen and Michael Hart for strains. Some strains were provided by the CGC, which is funded by NIH Office of Research Infrastructure Programs (P40 OD010440). This work was supported by NIH grant R01NS123645 (to P.T.K), a grant from the Mathers Foundation (to P.T.K.), and an NIH postdoctoral research award IRACDA-K12 GM102779 (to E.B.F).

## AUTHOR CONTRIBUTIONS

Conceptualization, E.B.F. and P.T.K.; methodology, E.B.F., A.T., Z.S., Y.W., and P.T.K.; software, A.T.; formal analysis, E.B.F. and A.T.; investigation, E.B.F., A.T., Z.S., P.S.H., and Y.W.; writing – original draft, E.B.F.; writing – review and editing, E.B.F, A.T., Y.W., and P.T.K.; visualization, E.B.F, A.T., Y.W., and P.S.H.; supervision, P.T.K.; project administration, E.B.F. and P.T.K.

## METHODS

### C. elegans methods

All *C. elegans* strains were derived from the Bristol strain N2 and raised at 20°C on NGM plates seeded with OP50 *E. coli*, according to standard protocols^91^. Animals were analyzed at the L4 larval stage (staged between L4.4-L4.6^92^) or as embryos as they developed from 1.5-fold through 3-fold stages. Transgenic strains were generated by microinjection, using 1-5 ng/μL of plasmid DNA injected alongside markers Podr-1::RFP (100 ng/μL) or Podr-1::GFP (100 ng/μL). A full strain list is included in Supplementary Table 1.

### Molecular biology

All plasmids were generated using Gibson isothermal assembly^93^ and are listed in Supplementary Table 2. Constructs used for expression in *C. elegans* were cloned into the pSM vector backbone^94^.

### Generation of CRISPR/Cas9-edited alleles

Endogenous gene edits were generated by gonadal injection of CRISPR-Cas9 protein complexes. These injection mixes contained 1.525 μM ALT-R Cas9, tracRNA, and guide RNA complexes (Integrated DNA Technologies) and either 6 μM ssDNA repair templates (ultramers from Integrated DNA technologies) or PCR-amplified repair templates injected at 1-5 μM, depending on the size/complexity of the edit. We employed a co-CRISPR strategy^95^, using either dpy-10 or unc-58, depending on the chromosome of the edit. Repair templates were designed with 100-200 bases of homology to each flank and were designed to either silently destroy the protospacer adjacent motif (PAM) site, or when silent mutations in the PAM were impossible, to silently mutate at least four nucleotides in the gRNA recognition sequence. All guide RNAs were synthesized as ALT-R CRISPR (cr)RNAs (Integrated DNA Technologies) and are listed in Supplementary Table 3. CRISPR-edited alleles were isolated through PCR screening, were verified by Sanger sequencing, and were outcrossed.

### Confocal microscopy

Confocal images of fluorescently labeled proteins were collected at room temperature in living, anesthetized *C. elegans* that were grown at 20°C. Approximately 20-30 L4-stage hermaphrodites were anesthetized in 20 mM levamisole (Sigma-Aldrich) in M9 buffer, mounted on 10% agarose pads, and sealed under a coverslip with Vaseline. Animals were staged again using DIC optics to exclusively image L4 animals within an ∼2-hour developmental window between L4.4-L4.6 stages^92^. Confocal z-stacks of the L4 dorsal nerve cord were collected with 0.27 μm z steps over a 6 μm z range. Confocal z-stacks of the L4 nerve ring were collected with 0.27 μm z steps over an 8 μm z range. ∼50 embryos per slide were mounted on a 10% agarose pad and imaged at stages indicated in figures (based on their morphology). Confocal z-stacks of embryos were collected with 0.3 μm z steps over a 6μm z range.

Images were acquired on a spinning disk system (3i) equipped with an Evolve 512 EMCCD camera (Photometrics) and a CSU-X1 M1 spinning disk (Yokogawa) mounted on an inverted Zeiss Axio Observer Z1 microscope. Endogenously labeled strains at all stages (L4s and embryos) were imaged with a Zeiss Plan-Apochromat 100x/1.4NA oil immersion objective. Strains expressing the DA9-specific marker wyIs685 were imaged with a Zeiss Plan-Apochromat 63x/1.4NA oil immersion objective. Images were acquired using Slidebook 6.0 software (3i).

### Confocal image processing and analysis

DA9 neuron and L4 dorsal nerve cord image analysis and visualization: Raw confocal z-stack volumes contained within the native .sld Slidebook file format (3i) were rendered into 16-bit maximum intensity projections tifs and 20-pixel lines were drawn over synaptic regions of interest and straightened into 32-bit tif linescans using Fiji^96^. Fluorescent puncta in the 32-bit linescan tifs were detected and segmented using a custom ROI-based MATLAB application (Image Processing Toolbox Release 2022a, The MathWorks, Inc.) that uses local means thresholding and ROI watershed segmentation followed by parametric restriction to remove noise pixels. The same thresholding and restriction parameters were applied to all images with particular fluorescent protein markers (e.g. NRX-1::Skylan-S). See Supp. Fig. 1 for examples of presynaptic puncta ROIs detected using this application. We calculated the number of puncta per 100 μm, the sum fluorescence intensity of puncta per 100 μm, and other ROI statistics that were not reported, including ROI area (um^2^) and ROI localization positions. Co-localization was measured as the % of NRX-1::Skylan ROI area co-localized with SYD-1 puncta. The linescans chosen for display in figures were most representative of the average calculated statistics for each genotype. The same linear adjustments to histogram parameters (minimum and maximum) were applied to all images displayed together across genotypes, and all images were displayed with inverted grey lookup tables (Fiji) for visualization.

L4 nerve ring image analysis and visualization: Raw data of confocal z-stack volumes contained within the native .sld Slidebook file format (3i) were rendered into 16-bit maximum intensity projections tifs. ROIs around the nerve rings were drawn using Freehand selections in Fiji and mean intensity was measured for each. Images that were most representative of the average calculated statistics were chosen for display, and the same linear adjustments to histogram parameters (minimum and maximum) were applied to all images shown together across genotypes. All images were displayed with inverted grey lookup tables (Fiji) for visualization.

Embryo image analysis and visualization: Raw confocal z-stack volumes in the native .sld Slidebook file format (3i) were rendered into 16-bit maximum intensity projection tifs in Fiji and assessed for the presence of endogenously-labeled synaptic proteins in the nerve ring at each stage. Embryos begin twitching just after the 1.5-fold stage, complicating our ability to collect clear, full z-stacks without movement artifacts and thus our ability to perform fluorescence intensity analysis of the embryonic nerve rings. Images that best displayed the nerve ring from each stage were selected from each z-stack (either as 2-3 image planes in maximum intensity projections or single image planes in cases where there were movement artifacts due to twitching). For visualization, embryo images were rotated to the same anterior/posterior and dorsal/ventral axes and were displayed with inverted grey lookup tables (Fiji) for visualization. We showed representative images of the developmentally earliest instances of detectable accumulation of SYD-1 or NRX-1, which we observed in at least five embryos across multiple imaging sessions, irrespective of fluorescent protein tag used. SYD-2/Liprin-α is detectable in embryos as early as the bean stage, as reported previously^23^.

### Atomistic molecular dynamics (AT-MD) simulations

AT-MD simulations were performed using the CHARMM36m force field^100^ in GROMACS 2023.1^101^. The AT-MD setup was generated using the CHARMM-GUI Bilayer builder^102,103^ which truncated SYD-1 to the domain of interest (PDZ, C2, or RhoGAP), generated the model membrane, positioned the protein above the membrane, and added in water extent. The system size was calculated based on a specific number of lipid molecules and an XY ratio of 1:1. We modeled membranes consisting of a total of 80 lipids, composed either entirely of POPC (control) or of 75% POPC: 15% POPS: 5% PIP_2_: 5% Cholesterol (PIP_2_-containing complete membrane). These membrane compositions resulted in XY dimensions of 7.39×7.39 nm^2^ for the control membrane and of 7.25×7.25nm^2^ for the complete membrane. The protein domain was positioned away from the membrane to ensure an initial separation greater than the Van Der Waals (VDW) and electrostatic cutoff distances of 1.2 Å. The Z dimension of the box was then calculated to guarantee a minimum water extent of 25Å at the top and bottom of the protein, resulting in a Z dimension of 1.52-1.73nm for the systems depending on the protein domain being simulated.

The system was first minimized and equilibrated for 2ns, and then 20 parallel production runs/trajectories of 200ns each were carried out. Both the equilibration and production runs used 2 fs time steps with 1.2 nm VDW and electrostatic cutoff distance. Equilibration was performed in the NVT and NPT ensemble and was run with an isotropic Berendsen barostat to maintain pressure at 1 bar with a time constant of 5 ps and water compressibility of 4.5⋅10^-5^bar^-1^ and with Berendsen temperature coupling^104^ applied separately to lipids, protein, and solvent, with a time constant of 1 ps to keep the temperature at 310 K. Production runs were performed using an isotropic Parinello-Rahman barostat^105^ to maintain pressure at 1 bar with a time constant of 5 ps and water compressibility of 4.5⋅10^-5^bar^-1^ and using Nose-Hoover temperature coupling^106^ applied separately to lipids, protein, and solvent, with a time constant of 0.5 ps to keep the temperature at 310 K.

Minimum protein-lipid distance was obtained using gmxmindist in GROMACS 2023.1 and runs where the minimum protein-lipid distance fell below 2Å were considered for further analysis. The PIP_2_ binding residues in the C2 domain were obtained from the SYD-1 C2/full membrane simulations using a custom tcl script in VMD^107^. Binding residues were defined as residues that stayed within 2Å of a PIP_2_ lipid for at least 50ns. Plots of minimum protein-lipid distance and binding residues were generated using R version 4.2.2^108–110^.

### Coarse-grained Langevin dynamics (CGLD) simulations

A coarse-grained model was designed to simulate the interactions between NRX-1 and SYD-1 in their cellular environments. This simplified model can reach the spatial-temporal scales that are not accessible for all-atom MD simulations but still maintains the features that are able to describe the basic dynamics of the system. Specifically, the system contains two proteins: NRX-1 and SYD-1. NRX-1 is further represented by two functional modules: a transmembrane (TM) domain and the PDZ binding motif (PBM). Each module is coarse-grained into a globular rigid body. The TM and PBM modules are further connected by a flexible linker. Similarly, SYD-1 is also represented by two functional modules, corresponding to the PDZ and C2 domains, respectively. Each domain is coarse-grained into a globular rigid body, and both domains are connected by a flexible linker.

Based on the model representation, an initial configuration was set up in which the plasma membrane is represented by the top surface of a three-dimensional cubic box and the cytoplasmic region is represented by the space below. Multiple copies of NRX-1 and SYD-1 polymers were then placed in the simulation box. The TM module of NRX-1 was positioned in the plasma membrane and was connected by its PBM module located in the cytoplasmic region. Both the PDZ and C2 modules of SYD-1 were initially distributed in the cytoplasmic region. The intramolecular conformation of each polymer was generated at random.

Following the initial configuration, the movements of all NRX-1 and SYD-1 polymers in the simulation box and their interactions were updated iteratively by our recently developed diffusion-reaction algorithm^111,112^. With each simulation step, the Brownian Dynamics (BD) algorithm was first applied to drive the diffusion of all proteins^113^. The diffusion of NRX-1 was confined because its TM region was constrained in the plasma membrane. On the other hand, both the PDZ and C2 domains of SYD-1 can freely diffuse in the cytoplasmic region. After diffusions were carried out for all proteins, the system checked the possibility that any new interaction might form between NRX-1’s PBM and SYD-1’s PDZ, depending on the distance between two specific modules and the association rate of the interaction. After association, a new intermolecular bond was formed, and the two proteins moved together. In the meantime, the system also checked for the possibility that any already-formed interactions might dissociate, depending on the corresponding dissociation rate. After dissociation, the intermolecular bond was removed, and two proteins moved separately. Additionally, if the C2 domain of SYD-1 moves close to the plasma membrane during simulations, it can be attached to the membrane surface, depending on the attach rate and detach rate.

The configuration of all proteins in the system and their interactions were updated after all above operations were completed for the current simulation time step. Through this repeated process, simulations were continued until equilibrium was reached.

### Prediction of the effect of radical amino acid replacement on protein stability

The effects of the C2* mutation (K629A; K633A; K635A; R692A; K694A; R725A) were assessed using the DDmut web server^49^. We obtained a pair of ΔΔG^stability^ values, one for the mutation prediction (WT->C2*, ΔΔG=0.45 kcal/mol) and one for the reverse prediction (C2*->WT, ΔΔG=-0.91 kcal/mol), showing that only slight destabilizing effects on the structure (Destabilizing ΔΔG are defined as >1 kcal/mol). We then evaluated the disruption of the C2 domain using VMD^107^. We calculated the backbone RMSD for the C2 domain in the native and C2* mutant, obtaining a value of 0.094 Å, well below the experimental variability of 1.2Å.

### Statistics

All data were presented as mean ± SD (standard deviation) on bar graphs unless otherwise noted, with individual data points displayed in scatter points overlaid on the bars. The sample size was defined as the number of worms and indicated at the base of each bar in all graphs. Datasets were collected over multiple imaging sessions over multiple days. Data were analyzed with Prism 10 (v10.0.2, GraphPad Software, Inc.). For comparison of two groups, unpaired T-tests with Welch’s correction were performed. For comparison of more than two groups, Brown-Forsythe and Welch ANOVA tests with Dunnett’s T3 multiple comparisons test were performed, unless otherwise noted. P<0.05 was considered significant.

## FIGURE LEGENDS

**Supplemental Figure 1.**
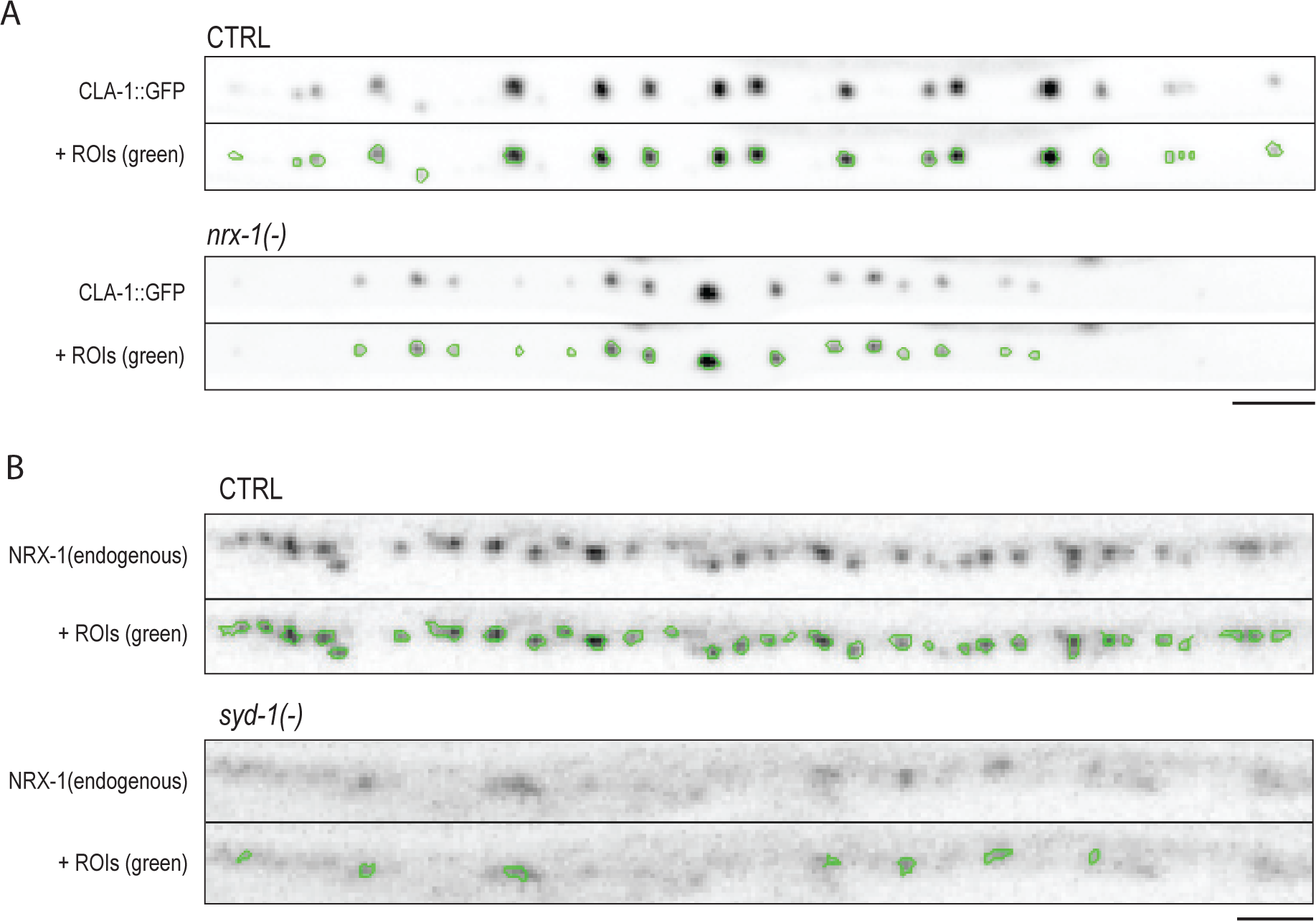
Example ROI identifications using custom MATLAB application. Supp 1A. Representative images of ROIs identified for DA9 neuron-specific CLA-1::GFP puncta in CTRL and *nrx-1(-)* strains. In each genotype, the top image is a representative linescan of CLA-1::GFP at presynapses, and the bottom image contains the ROIs (outlined in green) that were identified by our MATLAB script. The same thresholding and restriction parameters were applied to all images containing this marker. Scale, 5 μm. Supp 1B. Representative images of ROIs identified for an endogenous NRX-1::Skylan marker in a CTRL and *syd-1(-)* strain. The top image is a representative linescan of NRX-1::Skylan at synapses in the dorsal nerve cord, and the bottom image contains ROIs (outlined in green). The same thresholding and restriction parameters were applied to all images of the NRX-1::Skylan marker. Scale, 2.5 μm.

**Supplemental Figure 2.**
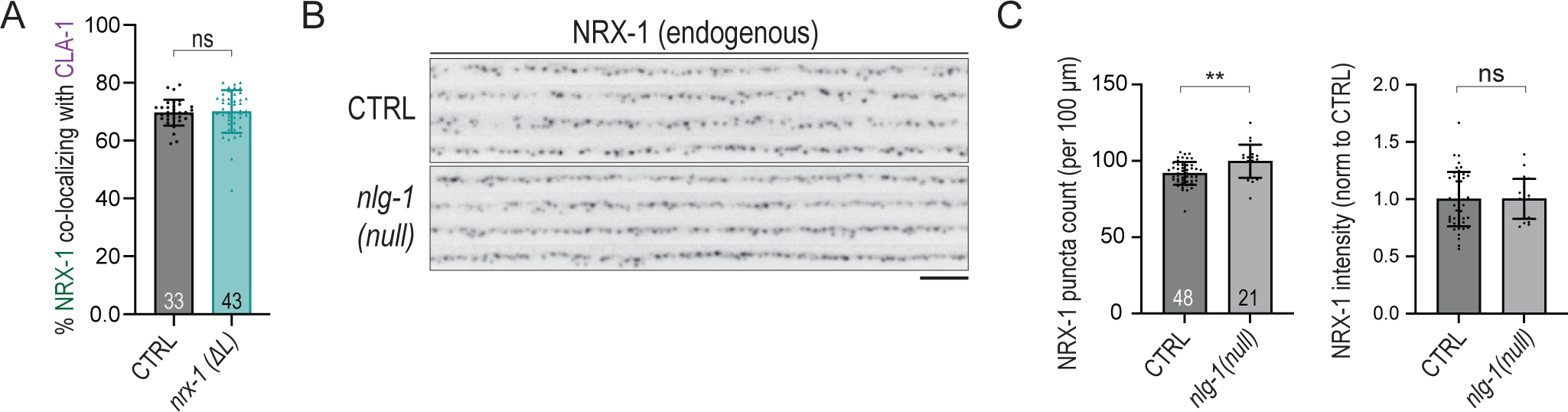
Neurexin does not depend on canonical binding partner neuroligin for synaptic localization. Supp 2A. Co-localization statistics of endogenous NRX-1::Skylan with CLA-1::mScarlet. Supp 2B. Representative images of endogenously labeled NRX-1::Skylan in the L4 dorsal nerve cord of *nlg-1(null)* mutants. Scale, 5 μm. Supp 2C. Quantification of NRX-1::Skylan puncta number and intensity in *nlg-1(null)*.

**Supplemental Figure 3.**
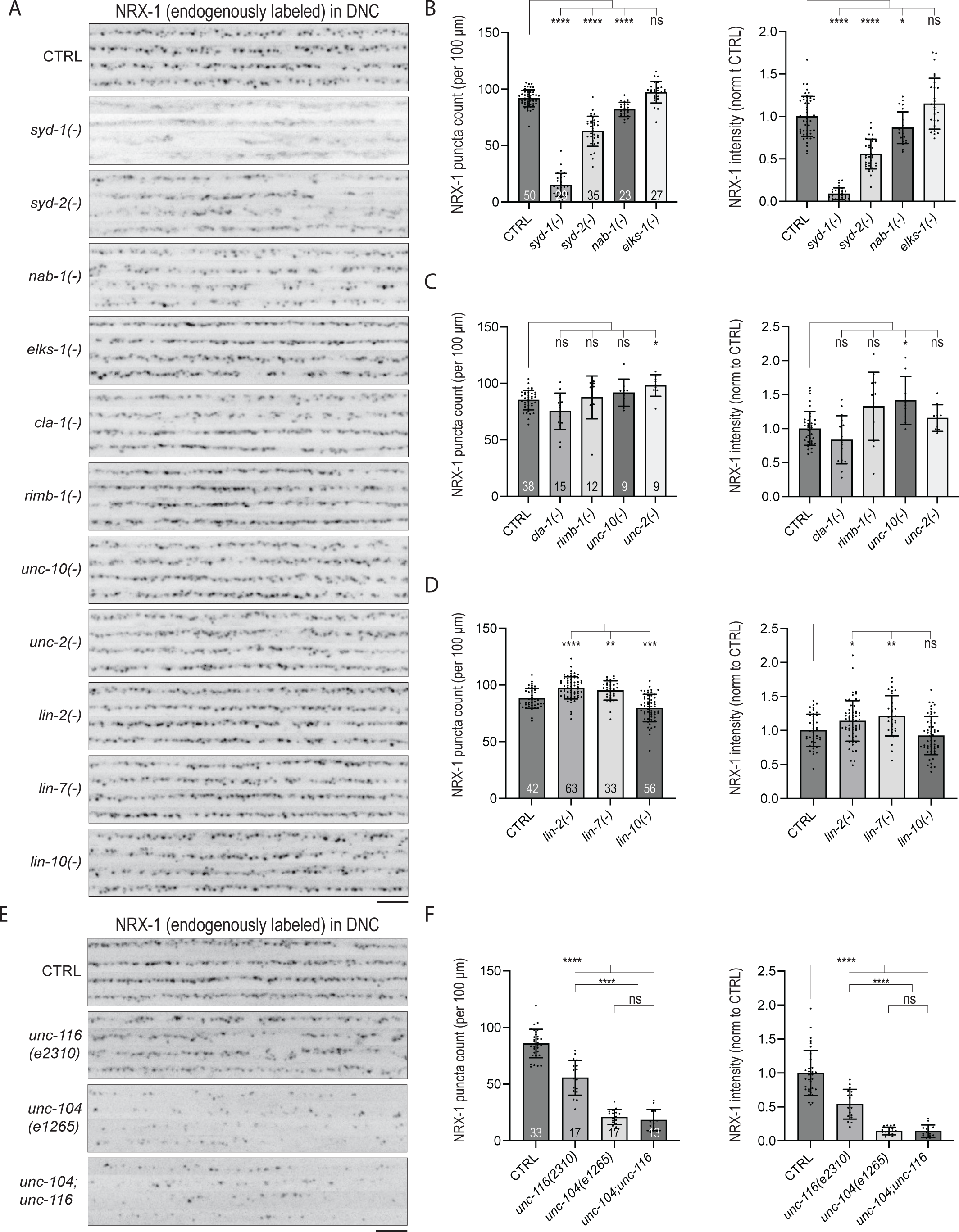
Candidate screen for intracellular proteins that regulate neurexin localization. Supp. 3A. Representative images of endogenously labeled NRX-1::Skylan in mutants of various presynaptic active zone genes. Scale, 5 μm. Supp 3B-D. Quantification of NRX-1 puncta number and intensity reveal severe disruption in *syd-1* mutants, as well as significant defects in *syd-2* and *nab-1* mutants. Mutants of *elks-1*, *cla-1,* and *unc-2* did not appear to lead to any defect in NRX-1 puncta number or intensity. Mutants of *rimb-1/*RIM-binding protein and *unc-10/*RIM led to a slightly elevated NRX-1 puncta intensities. Mutants of proteins predicted to interact with the intracellular domain of mammalian neurexin *lin-2/*CASK*, lin-7*/Veli, and *lin-10/MINT* displayed mild but significant differences in NRX-1 distribution. Supp 3E. Representative images of NRX-1::Skylan in kinesin-1 *unc-116/*KIF5A/C and kinesin-3 *unc-104/*KIF1A mutants. NRX-1 trafficking is predominantly mediated by the Kinesin-3 protein UNC-104/KIF1A, with a minor contribution from UNC-116, consistent with previous observations^32^. Note that the NRX-1 that localizes to axons in the absence of kinesin trafficking machinery is still able to cluster. Both *unc-104(e1265)* and *unc-116(e2310)* are partial loss-of-function/hypomorphic alleles. Scale, 5 μm. Supp 3F. NRX-1 puncta number and intensity are significantly reduced in both *unc-116* and *unc-104* mutants but show a major reliance on *unc-104.* The double mutants of *unc-104(e1265);unc-116(e2310)* are not more severe than the *unc-104(e1265)* mutants.

**Supplemental Figure 4.**
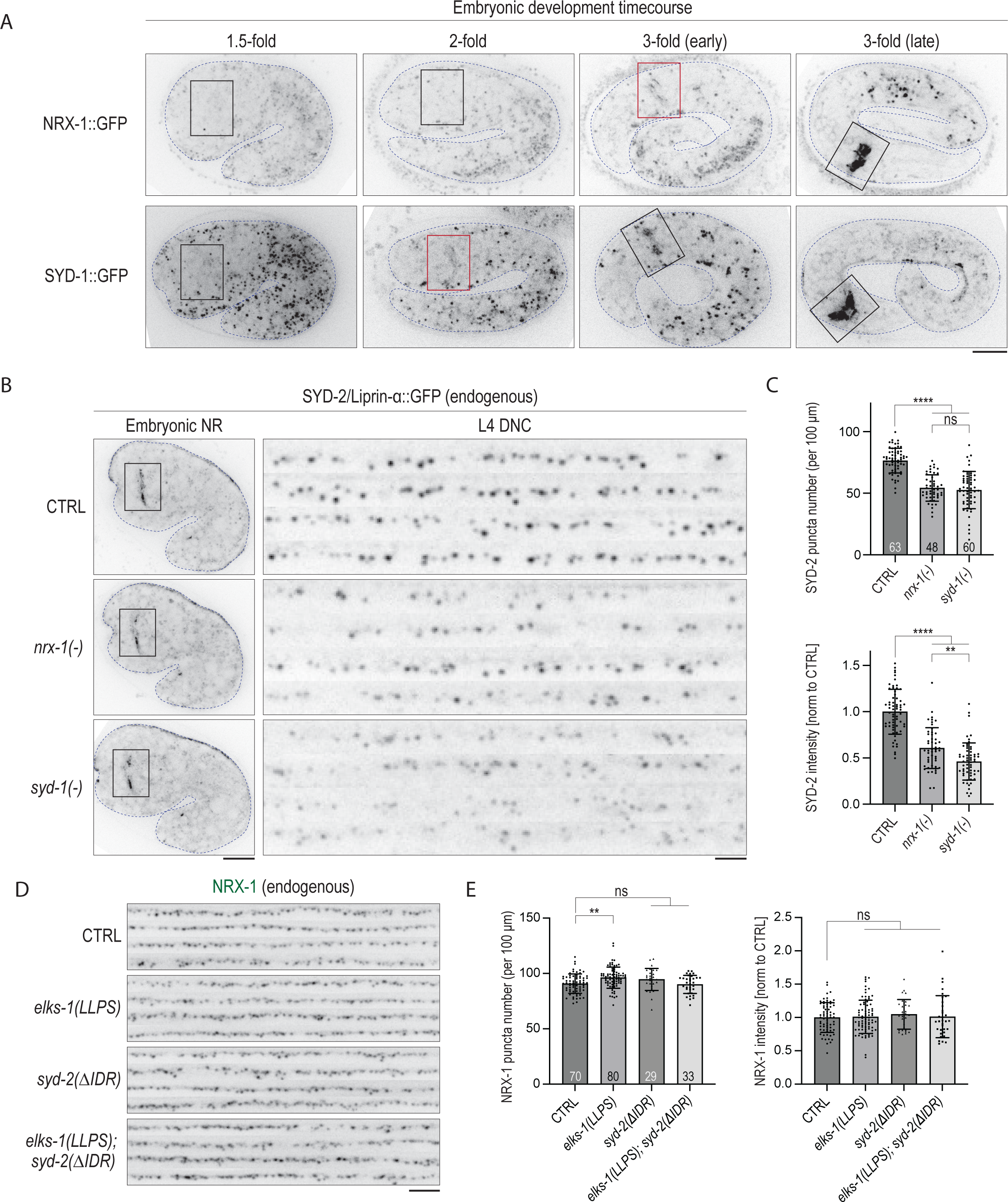
NRX-1 functions in stabilization, but not initial accumulation, of SYD-2 at presynapses and does not require phase separation for its own localization. Supp 4A. Endogenously labeled NRX-1::superfolderGFP or SYD-1::GFPnovo2 display the same expression patterns in embryos as those displayed in 4A, when each protein was labeled with a different fluorescent protein. SYD-1 is first detectable in the embryonic 2-fold stage nerve ring and NRX-1 is first detectable in the 3-fold stage nerve ring (red boxes). Scale, 10 μm. Supp 4B. SYD-2/Liprin-α accumulation in embryos is not affected by mutants of *nrx-1* or *syd-1,* but its long-term retention in presynaptic puncta is reduced in mutants of both genes. Embryo scale, 10 μm. DNC, 2.5 μm. Supp 4C. Quantification of SYD-2 puncta number and intensity in L4 DNCs. Supp 4D. Representative images of endogenously-labeled NRX-1::Skylan in mutants that disrupt liquid-liquid phase separation properties of SYD-2 and ELKS-1. These mutants have no obviously distinguishable impact on NRX-1 localization or clustering in the L4 dorsal nerve cord. Scale, 5 μm. Supp 4E. NRX-1 puncta number (left) and intensity (right) are not reduced in either of the phase separation mutants, or a double mutant containing both mutations.

**Supplemental Figure 5.**
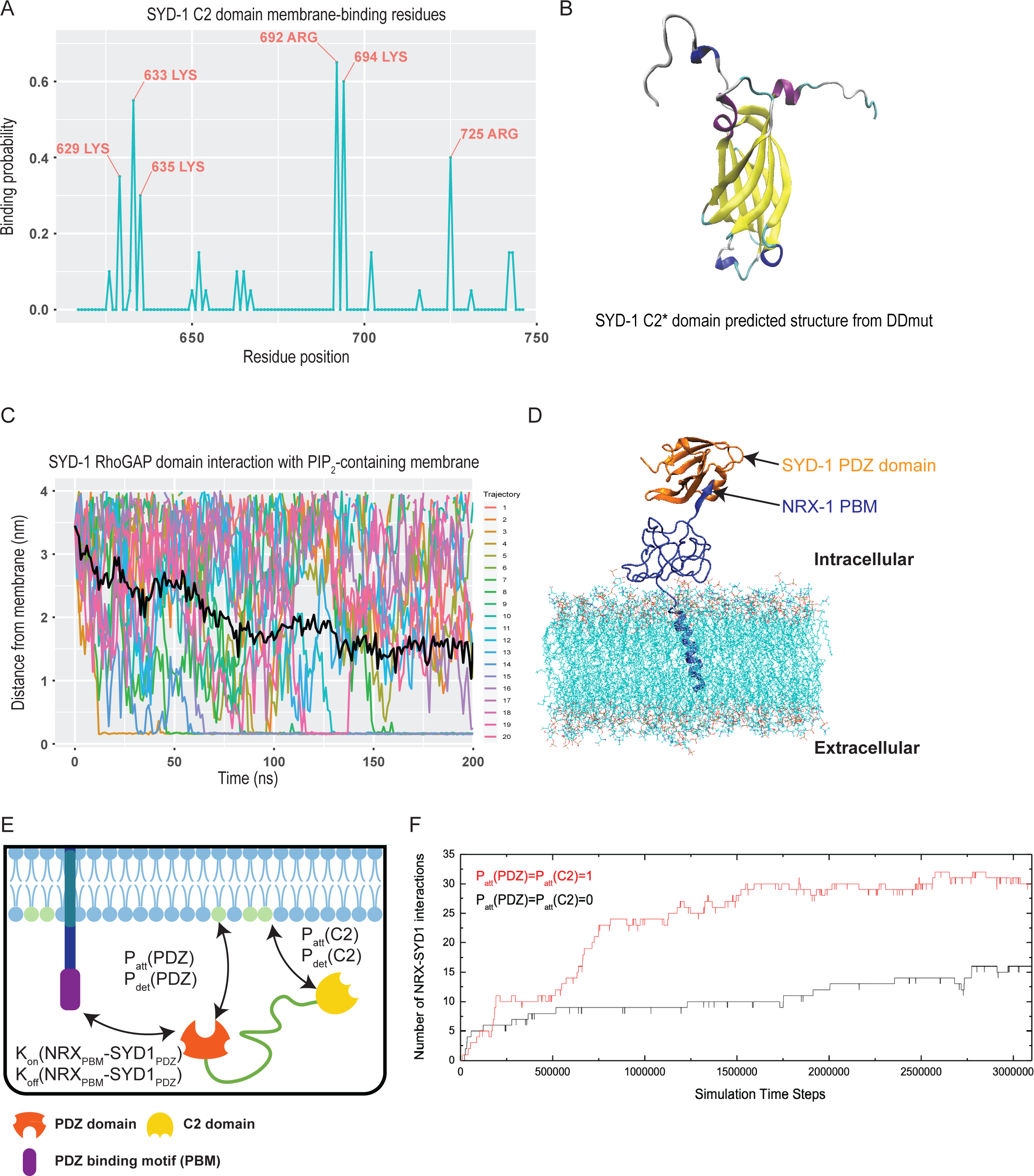
Interaction of SYD-1 PDZ and C2 domains with membrane increase the probability of PDZ-based interactions between SYD-1 and NRX-1. Supp 5A. Probability of PIP_2_ binding (>50 ns of binding within the 200ns simulation) for each residue of the SYD-1 C2 domain based on 19 trajectories. Labeled amino acids have 30% or greater probability of binding PIP_2_, suggesting that these are the residues that facilitate the SYD-1 C2 domain’s interaction with membrane. Supp 5B. AlphaFold 2.0-predicted structure of SYD-1 C2 modified by DDmut to contain substitutions of PIP_2_-interacting amino acids identified in 5A for alanine. The SYD-1 C2* domain predicted secondary structure is very similar to the native SYD-1 C2 domain, with an average ΔΔG^stability^ value of 0.68 kcal/mol and RMSD of 0.094Å. Supp 5C. Unlike the SYD-1 C2 and PDZ domains, the SYD-1 RhoGAP domain was not predicted in MD simulations to engage in stable interactions with PIP_2_-containing membrane within 200 nanoseconds. Plots depict the minimum distance between the SYD-1 RhoGAP domain and the same theoretical membrane used in MD simulations of the C2 and PDZ domains (75% POPC, 15% POPS, 5% PIP_2_, 5% cholesterol). The black line represents the rolling average distance between the domain and membrane at each time point. Supp 5D. Simulated interaction between the SYD-1 PDZ domain and the NRX-1 PDZ binding motif (PBM) at the plasma membrane. Supp 5E. Course-grained Langevin dynamics (CGLD) simulation predicts how the SYD-1 PDZ and NRX-1 PBM interaction is affected by the attachment of the SYD-1 PDZ and C2 domains to the membrane. Supp 5F. Plots of the number of predicted interactions occurring between SYD-1 PDZ and NRX-1 PBM under conditions in which the SYD-1 PDZ and C2 domains are stably interacting with the membrane (blue), or not interacting with the membrane (red). The interactions between the SYD-1 PDZ domain and NRX-1 PBM motif are generally increased as the attachment of the SYD-1 PDZ and C2 domains to the membrane are increased.

**Supplemental Fig 6.**
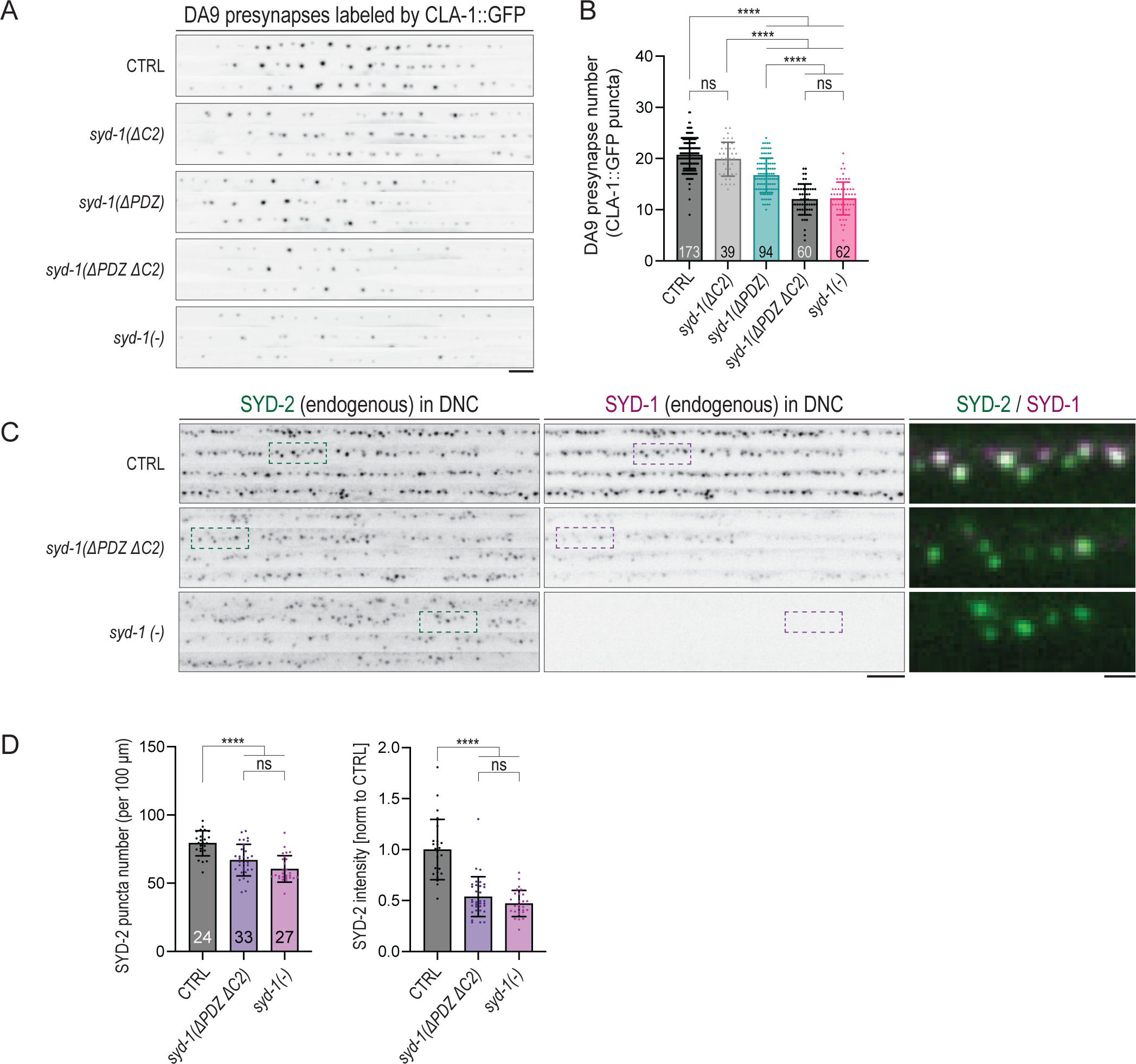
The C2 and PDZ domains of SYD-1 redundantly mediate synaptic assembly/stabilization *in vivo*. Supp 6A. Representative images displaying the effects of *syd-1* mutants on DA9 neuron synapses. Scale, 5 μm. Supp 6B. Quantification of *syd-1* mutant effects on DA9 presynapse number. *syd-1(ΔPDZ ΔC2)* and *syd-1(-)* mutants have similar effects on synapse number, while the loss of either the PDZ or C2 domains alone display relatively minor (or no) reductions in synapse number. Supp 6C. Representative images of endogenously labeled SYD-2::GFP and SYD-1::mScarlet in *syd-1* mutants. Scale, 5 μm. Merge scale, 1 μm. Supp 6D. Quantification of SYD-2 puncta number (left) and intensity (right) in *syd-1* mutants.

---

**Supplemental Table 1.**
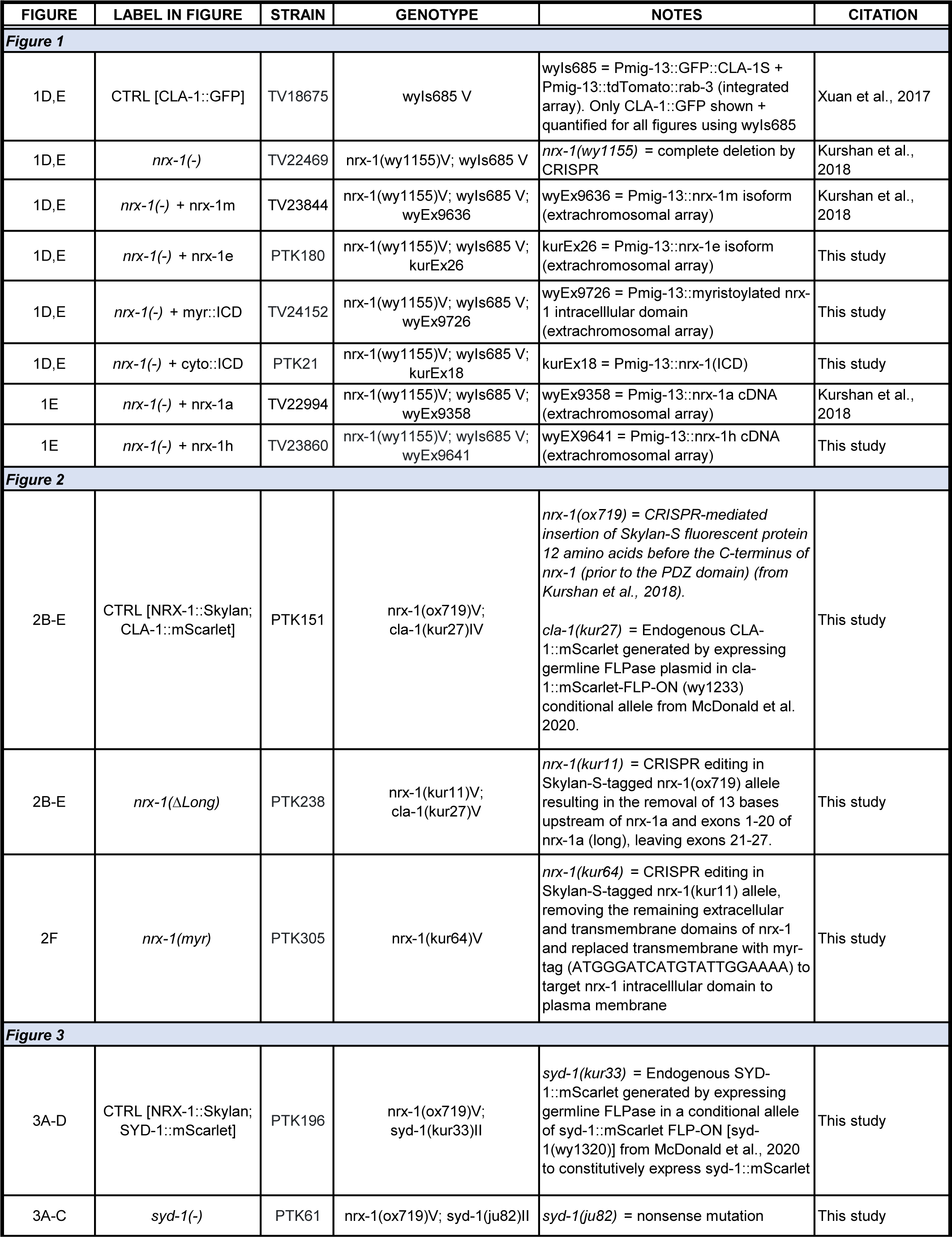
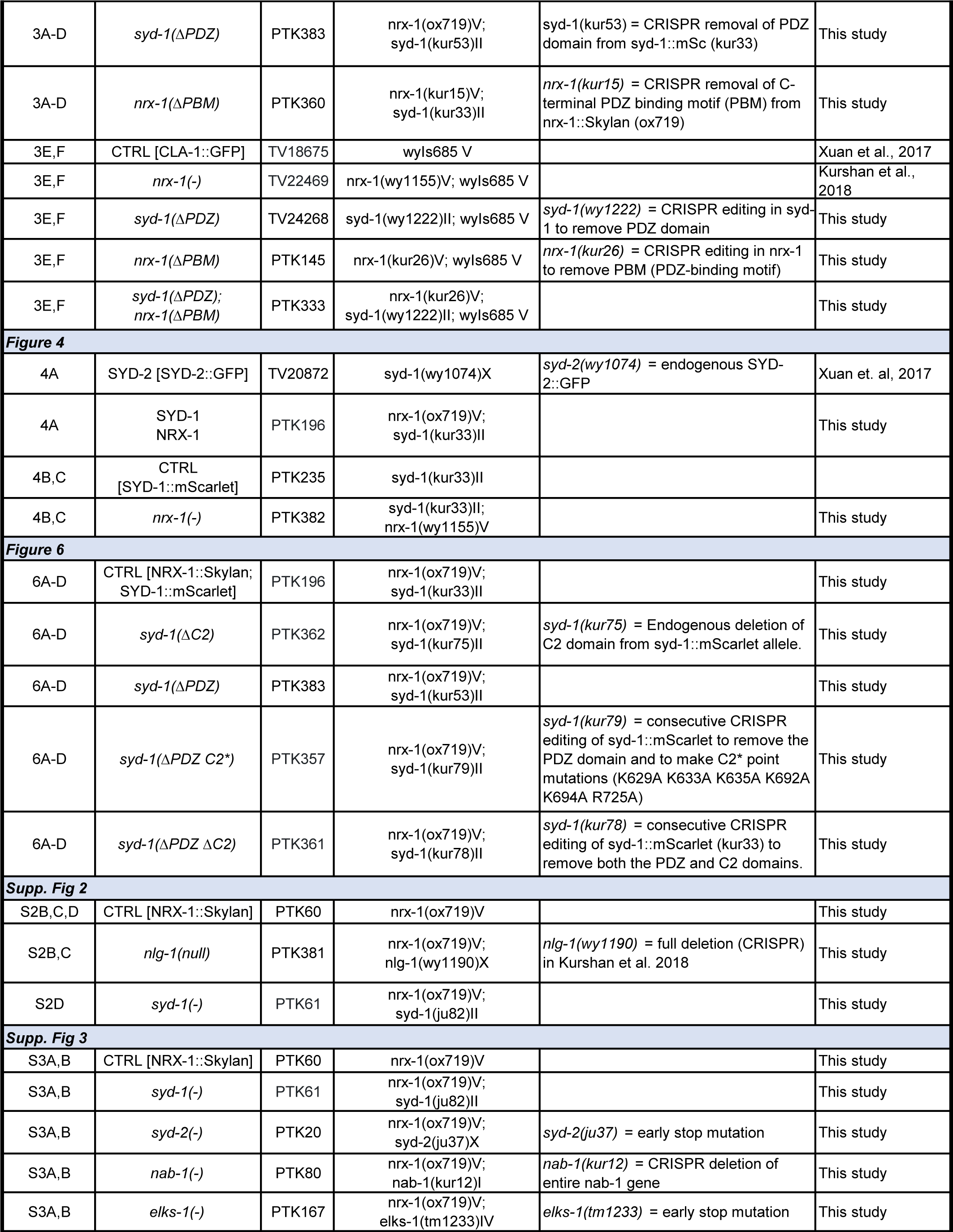
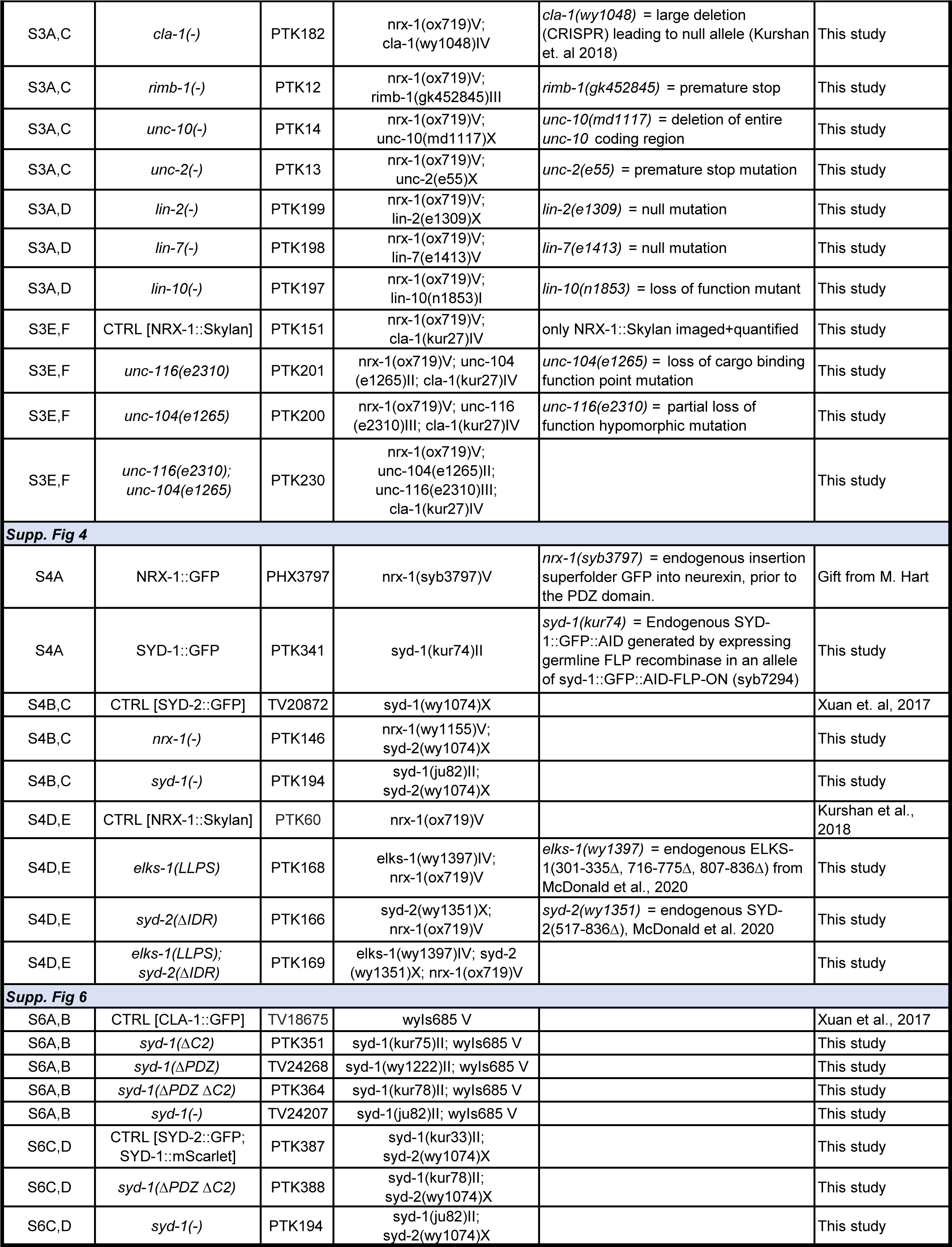

**Supplemental Table 2.**
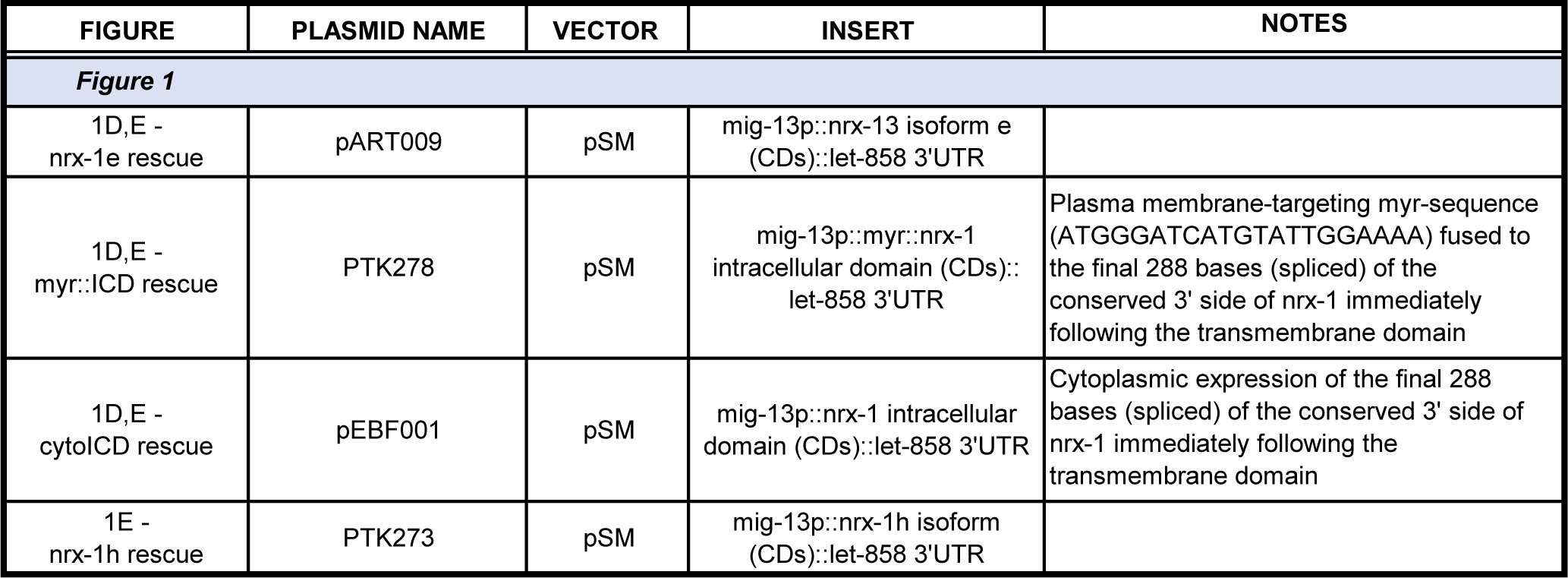

**Supplemental Table 3.**
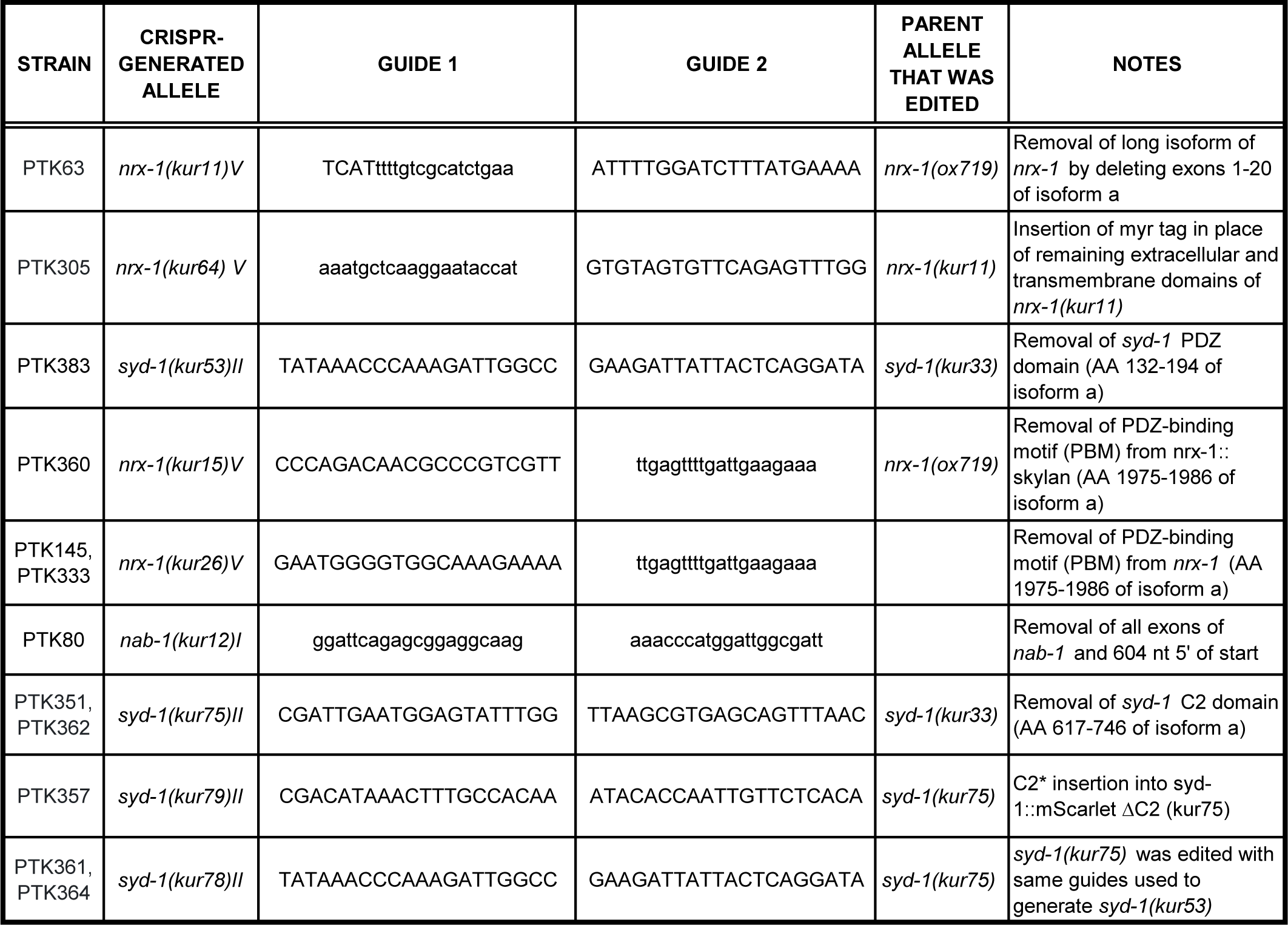

